# Microbial coexistence through chemical-mediated interactions

**DOI:** 10.1101/358481

**Authors:** Lori Niehaus, Ian Boland, Minghao Liu, Kevin Chen, David Fu, Catherine Henckel, Kaitlin Chaung, Suyen Espinoza Miranda, Samantha Dyckman, Matthew Crum, Sandra Dedrick, Wenying Shou, Babak Momeni

**Affiliations:** Department of Biology, Boston College, Chestnut Hill, MA 02467; Department of Computer Science, Boston College, Chestnut Hill, MA 02467; Division of Basic Sciences, Fred Hutchinson Cancer Research Center, Seattle, WA 98109

## Abstract

Many microbial functions happen within communities of interacting species. Explaining how species with intrinsically disparate fitness can coexist is important for applications such as manipulating host-associated microbiota or engineering industrial communities. Previous coexistence studies have often neglected interaction mechanisms. Here, we formulate and experimentally constrain a model in which chemical mediators of microbial interactions (e.g. metabolites or waste-products) are explicitly incorporated. We construct many instances of coexistence by simulating community assembly through enrichment and ask how species interactions can explain coexistence. We show that growth-facilitating influences between members are favored in assembled communities. Among negative influences, self-restraint, such as production of self-inhibiting waste, contributes to coexistence, whereas inhibition of other species disrupts coexistence. Coexistence is also favored when interactions are mediated by *depletable* chemicals that get consumed or degraded, rather than by *reusable* chemicals that are unaffected by recipients. Our model creates null predictions for coexistence driven by chemical-mediated interactions.

## 1. Introduction

Microbial communities influence ecosystems by cycling matter (1) and affect human health by facilitating food digestion or causing infections (2–5). Cohabiting microbes in communities interact and can perform functions that none of the member species can achieve efficiently on their own (6, 7). Examples of such functions include degradation of complex compounds such as crude oil, cellulose, or plastics (8–10). How can species in a community stably coexist, despite differences in their intrinsic fitness? To design long-lasting communities for waste remedy or fuel production (11, 12), or to manipulate host-associated communities (13, 14), a better understanding of coexistence mechanisms will be instrumental.

Exploring the mechanisms that would allow coexistence (defined as extended presence of different species within a community) and stability (defined as maintenance of coexistence despite perturbations) has been among major directions in community ecology (15–19). Many studies, both theoretical and experimental, have identified how coexistence may be achieved. Trade-off in life traits between colonizers and competitors (20), refuges for prey in a spatially structured environment (21, 22), and frequency-dependent fitness in mutualism (23–25) are a few examples of mechanisms identified in simple communities. Such coexistence mechanisms can be split into two broad categories: neutral theory or niche theory (26). Simply put, neutral theory deals with situations where either species have similar competitive abilities or other processes such as migration define the community. Interactions play a minor role within this framework. In contrast, niche theory deals with situations where interactions (either among species or between species and their environment) are strong and major drivers of coexistence. While depending on the situation one or the other framework would be more suitable for explaining coexistence, here we will focus on the impact of interspecies interactions (within niche theory) on coexistence.

Species interactions have been the subject of previous coexistence studies. Facilitation (i.e. interactions that benefit at least one of the partners), for example, has been identified to support coexistence. Facilitation may contribute to coexistence in different ways: boosting the growth of intrinsically less fit recipients (27, 28), relieving facilitators from competitive pressure (27), or protecting vulnerable species from harsh environments or predators (28, 29). Identifying other mechanisms of coexistence will expand our understanding of how communities form and persist.

To identify other mechanisms of coexistence, experimental studies in complex natural communities can be challenging. An alternative path is to use mathematical modeling. Compared to studies in natural communities that often involve a plethora of poorly characterized interactions, mathematical models offer a transparent and controllable platform (30). Modeling allows investigating conditions that are not easily attainable experimentally, e.g. by allowing full control over interaction parameters (31). Additionally, modeling enables screening many scenarios, a fate practically not feasible in experiments.

For studies in mathematical models, an important decision is how to represent the communities (32). Different possible mathematical representations can be categorized into three groups (33): 1) “association” models that solely rely on which species interacts with each of the other species, 2) “fitness” models that quantitatively include how species influence the fitness of each other, and 3) “mechanistic” models that include the mechanisms of interactions.

Most previous models of communities abstract all the interactions between species into pairwise fitness interactions (34–38). These models are intended to recapitulate how each interaction influences the fitness of the two involved parties (15, 35, 34, 14). Wide-spread use of pairwise fitness models presumably has two roots. First, from a historical perspective, many early community studies were built on prey-predation food webs (17, 39, 40) or plant-pollinator mutualisms (41–43) that relied on direct encounters without interaction mediating agent. Second, from a technical perspective, quantifying the interaction mediators, such as concentrations of exchanged metabolites, has been very challenging. The simplifying assumption of pairwise fitness models allows modeling the community without the challenge of characterizing the mediators (44). However, pairwise fitness models may not accurately capture common situations in which interactions take place through different mechanisms (e.g. by depletable or reusable mediators), when multiple influences take place through different mediators, or when shared mediator are produced or consumed by multiple species (44).

To overcome the limitations of pairwise fitness models, here we use a mechanistic model framework which incorporates chemical mediators of interspecies interactions (44). This choice is motivated by three factors: 1) indirect interactions where one species affects how strongly other species interact (45–48) are lost in pairwise interaction models but preserved in mechanistic models. 2) Interactions mediated by chemicals (e.g. metabolites or toxins) are common among microbes (5, 49, 50), and are thought to play a major role in microbial communities. 3) Recent progress in techniques such as stable isotope probing (SIP) (51), mass spectroscopy (MS) (52, 53), and nuclear magnetic resonance (NMR) (54) has improved our ability to identify and quantify interaction mediators. Thus, obtaining experimental data to constrain these mechanistic models is possible.

To examine the mechanistic origins of coexistence, we examine instances of simulated coexistence and ask what features of their interaction network may contribute to coexistence. To find instances of coexistence, we simulate a typical enrichment process to assemble communities. Computationally, we start from an initial pool with many species engaged in randomly assigned chemical-mediated interactions. We simulate how these species interact and grow in an environment with the shared resource in excess (similar to a turbidostat). Often, some of these species go extinct and a subset of species (one or more) persist within a span of tens of generations. If there are more than one species in the enriched community, we consider it an instance of species coexistence. We repeat this process of enrichment many times, each time starting from an initial pool of species with randomly assigned interactions, compiling an ensemble of instances of coexistence in communities.

Within the ensemble of coexistent communities, we search for common features. As discussed below, our results show that interaction can indeed drive coexistence in multispecies communities. We observe that this coexistence depends on how interactions take place (e.g. through a signaling molecule or a consumable metabolite), not just their fitness effect. As a result, we show that only under specific conditions, when interactions are mediated through independent, reusable mediators, the commonly used fitness models can accurately predict coexistence. We see that facilitation (i.e. stimulation of growth of other community members) is favored in coexistent communities, whereas inhibition of other species (but not self) is disfavored. We also observe that in many instances, these effects are causal; in other words, facilitation and self-restraint (i.e. inhibition of self) interactions encourage coexistence, but inhibitory interactions that suppress other species are detrimental to coexistence.

## 2. Results

### 2.1. Mechanistic model of chemical-mediated interactions

We model communities in which each species interact with other members of the community through chemical mediators (Fig 1A) (44). Each species can produce multiple chemicals and each chemical can influence multiple species (Fig 1B). To clarify our nomenclature, “interactions” are how species impact the fitness of their own type (intraspecies) or other species (interspecies). Each of these interactions might be the result of multiple “chemical influences,” or “influences” for short. Each influence in our model represents how a chemical produced by community members affects the fitness of a certain species. In our network, we indicate a relation from chemical to species as “f-link,” if there is a fitness influence by a chemical on a species, and a relation from species to chemical as “c-link”, if a species changes the concentration of a chemical (e.g. by production or consumption). A “link” may refer to a c-link or an f-link. Even though microbes are expected to change many chemicals in their environment (53, 55), in our model we limit the number of mediators. Our reasoning is that (1) we are only including mediators with strong fitness influence, and (2) recent studies suggest that different mediators can be grouped into “functional” categories, such as organic acids or small sugars, simplifying community models (56).

**Fig 1.**
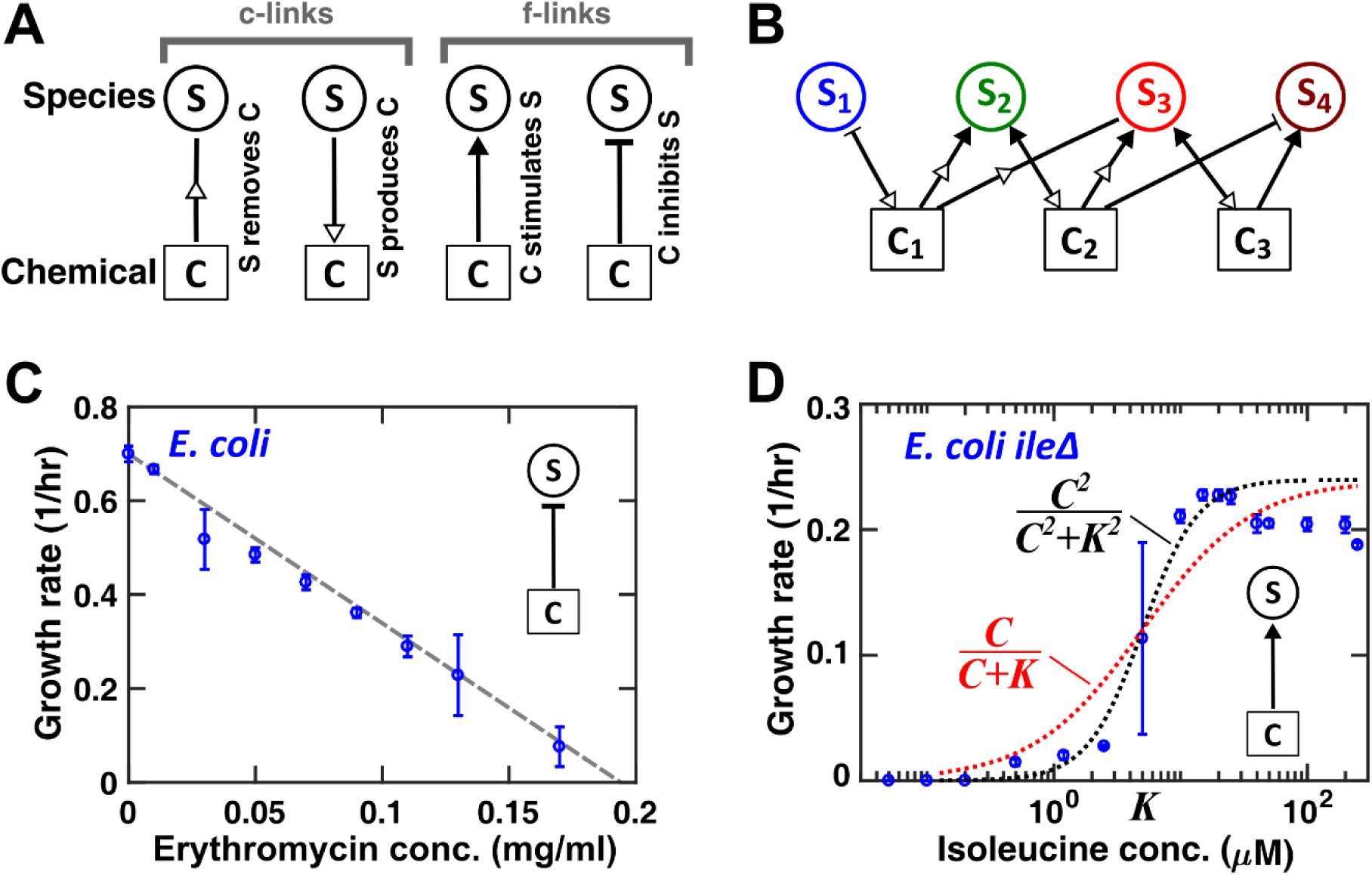
By explicitly incorporating the chemicals, we simulate a community of microbes engaged in chemical-mediated interactions. (A) Community networks are defined by two types of links: c-links (chemical production/removal links) indicated by hollow arrowheads and f-links (fitness influences on species) indicated either by filled arrows for facilitation or bar-termination for inhibition. (B) A combination of c-links and f-links can represent a community of species interacting through chemical mediators. (C) The influence of an inhibitory chemical on species is assumed to linearly affect the growth rate, motivated by experimental measurements of growth in presence of antibiotics (*E. coli* K-12 is used here as a representative example). A similar trend is observed in inhibitory effect of metabolic byproducts (Fig 1-FS1) on several strains. (D) The influence of a facilitative chemical compound on species is approximated to follow the Monod’s equation for simplicity. Experimental observations of auxotrophic *E. coli* K-12 strains suggest that a second-order relation (black dotted line) offers a more accurate estimation, but the first-order Monod-type equation (red dotted curve) is still an acceptable approximation.

To build realistic assumptions into our model, we assessed examples of how chemical mediators could affect the growth of cells. We experimentally characterized growth of bacterial cells in the presence of different concentrations of chemical compounds, C_*l*_, that stimulated or inhibited growth (Fig 1C-D). For growth inhibitors (Fig 1C), we have frequently observed that the growth rate linearly drops as the concentration of the inhibitor increases (Fig 1-FS1). For inhibition by antibiotics, we have observed that the inhibition is typically exerted above a threshold concentration (Fig 1-FS1), but for simplicity, in our model here we assume that threshold to be negligibly small for all inhibiting chemical influences. This model is slightly different in form compared to a previous report (57) that suggested a “threshold” inhibition effect, but we do not expect this difference to impact the overall outcome of our investigations. For growth facilitators, we observe the common biological situation in which over-abundance of the mediator does not proportionally contribute to the fitness effect (Fig 1D). Our work (Fig 1D; Fig 1-FS2) and others’ (57) suggest that a general saturating form, Moser equation 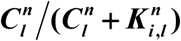, provides a good approximation for many cases of response to growth facilitators. ***K _i,l_*** is the concentration that parametrizes the saturating form of the dependence on the chemical concentration. For simplicity, we adopt the simplified Monod form ***C_l_/(C_l_+ K_i,l_***) to model this saturating behavior. We also assume that consumption is proportional to growth rate, with the same saturating form ***C_l_/(C_l_ + K_il_*)**. This is motivated by the assumption that acquisition of metabolites is the main factor impacting the fitness of cells.

By representing chemical concentrations as ***C_1_***, …, ***C_M_*** and live species abundances as ***S_1_***, …, ***S_N_***, changes in concentrations of chemicals and populations of species can be described in our modified model as

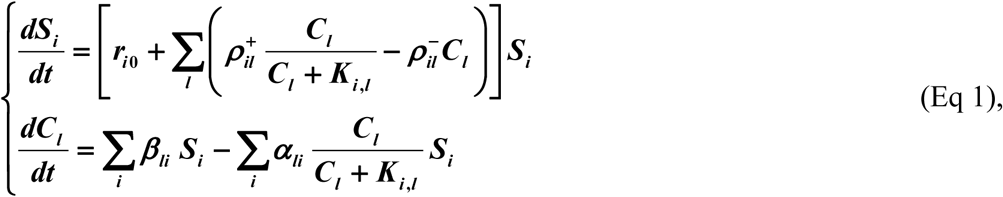

where *α_ik_* is the rate of consumption of ***C_i_*** per ***S_k_*** cell, ***β_ij_*** is the rate of production of ***C_i_*** per ***S_j_*** cell, ***r_i0_*** is the net basal growth rate of *S_i_* in the absence of chemical-mediated interactions, and ***ρ_il_*** represents the (positive or negative) influence of *C_l_* on the growth rate of ***S_i_***. Note that the death rate in this formulation is absorbed into the net growth rate, ***r_i0_***, and only live cells (with concentrations represented as ***S_i_***) contribute to removal and production of chemicals.

We assume in our model that each producer population can produce or consume multiple mediators, and each mediator can stimulate or inhibit the growth of recipient species (Fig 1A-B). Lastly, the combined effect of multiple mediators on the growth rate of each recipient species is assumed to be additive, similar to McArthur’s model of resource utilization (58). This assumption may capture, for example, the situation that a species can grow on multiple metabolites (59), but not the situation in which more than one mediator is necessary for the growth of a species. Previous reports have also discussed the exact form that may be more appropriate for representing the combined fitness influence of combinations of certain carbon sources (60, 61). For simplicity, here we do not incorporate this level of detail for modeling the influence of chemical mediators and use the additivity assumption (Eq 1) as an approximation. To focus on intercellular interactions, we assume that shared resources do not become limiting, although the model can be easily adjusted to incorporate shared resources (44).

### 2.2. Enrichment process can lead to communities that exhibit coexistence

We simulate enrichment for assembling coexistence. Enrichment, in the environmental or laboratory setting, is referred to the process in which supplying resources to an initial assembly of microbes affect the future population structure (62–64). During enrichment, starting from many initially interacting species, a coexisting subset can emerge (65). We will simulate enrichment *in-silico* (similar to (35)) by starting from several interacting species in an initial pool and following their dynamics over time. As the populations grow amidst these interactions, some species are outcompeted, whereas others coexist (Fig 2A, bottom; see Methods-Implementation).

**Fig 2.**
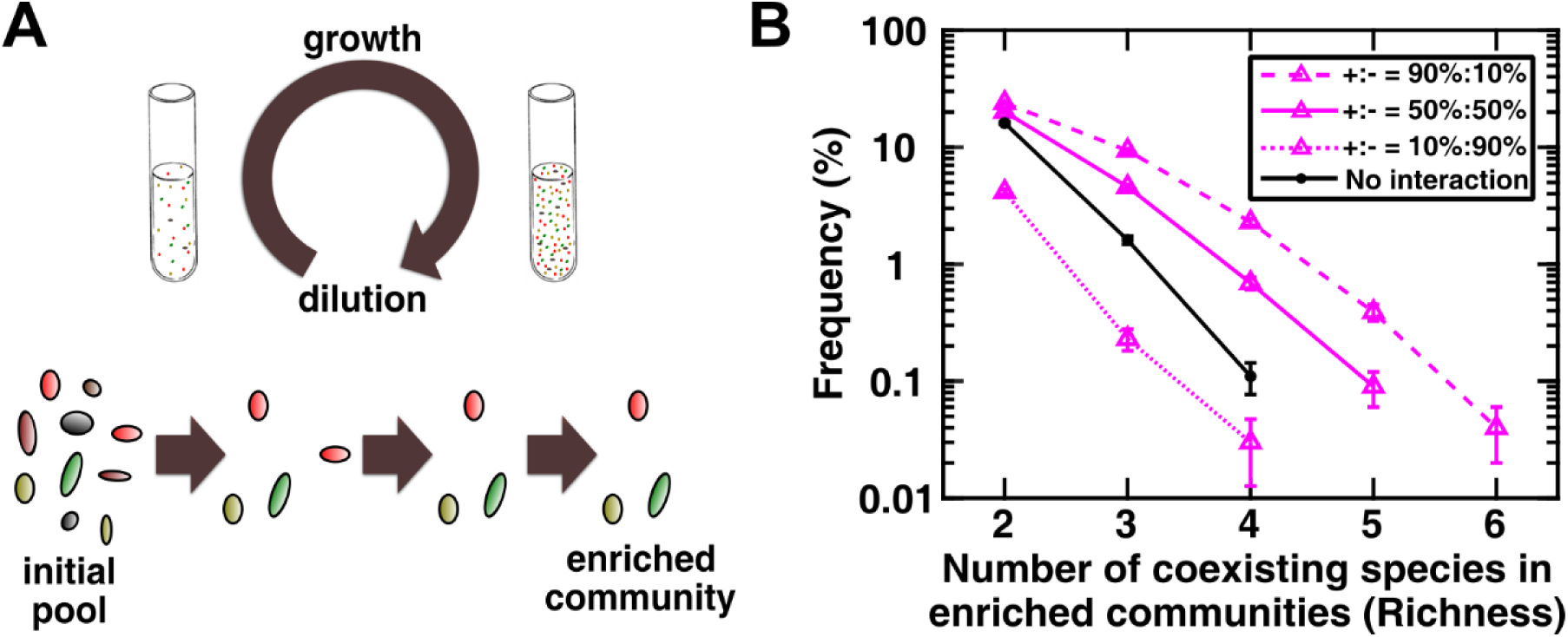
Simulated enrichment produces instances of species coexistence. (A) We simulate a typical experimental process of enrichment, in which a community of several species is grown in excess shared resource and are periodically diluted to a set density. In this scheme, we remove species that drop below a viable threshold abundance (corresponding to 1 cell). After several rounds of dilution (around 100 generations), we have observed that a subset of species remain in the community. In most cases, these species stably coexist if the dilution scheme is further extended to 200 generations. We take communities that contain more than one species at 150 generations as instances of communities that exhibit coexistence. (B) The likelihood of coexistence declines with community richness and increases with facilitative interactions. Community richness is defined as the number of coexisting species. As a reference, we have calculated the likelihood that for the same set of parameters, in the absence of any interactions we would observe coexistence (solid black curve). In these simulations, we have assumed that chemicals that influence a species, in half of the cases are depletable (***αli*** = 0 in Eq. 1), whereas in the other half, they are reusable (***αli*** = 0 in Eq. 1). We chose the initial number of species types N_c_=20 and the number of possible mediators N_m_=15.

In each simulation, we initially put together several species with a random network of interactions at equal proportions. Assuming a well-mixed coculture, these cocultures grow from a set initial population size up to a dilution threshold level, upon which the culture is diluted back to the initial population size (Fig 2A, top). This process reflects a common experimental practice where shared resources (including space) are replenished. As a result, the inherent assumption in our simulations is that cells are not limited by any of the shared environmental resources. Instead, mediators produced by cells determine community interactions and thereby coexistence. This allows us to focus primarily on how intercellular interactions can contribute to species coexistence. Growth is simulated for 15 dilution cycles which is typically enough to reach a stably repeating pattern of population dynamics within each cycle. In this process, species that drop below a population size of one cell are considered extinct and removed from the rest of the simulation (Fig 2-FS1). The remaining species are considered to coexist (Fig 2A, bottom), within the time-scale of our run, and we record the resulting community as an instance of coexistence. Note that coexistence in this case is functionally defined as maintenance of species together during our enrichment process, from an experimentalist’s point-of-view (for example, as in (66)).

Figure 2B shows the likelihood of reaching communities of different richness, starting from a pool of randomly interacting species. In these simulations, we have assumed that the initial pool of species has a random connectivity network in which c-links and f-links each have a fixed presence-absence probability (i.e., each have a Erdős–Rényi connectivity graph (67); see Methods-Implementation). We will call such networks “binomial” throughout this work. Under the assumptions of this simulation, listed in Supplementary Information (Simulation parameters), the model predicts that the likelihood of achieving communities with higher species richness, starting from an initial pool of randomly interacting species, decreases at an exponential rate. As a control, we examined a community with similar parameters, but with no interactions (representing the situation within the neutral theory of coexistence). In this situation, species with highest fitness can coexist, if their basal fitness happens to be close enough. The chance of coexistence also exponentially drops for communities with higher richness (Fig 2B, black curve). In our model, since each species can independently grow with a basal fitness in the enrichment environment, we can assume that the number of resources (as typically defined in a consumer-resource model) is at least as many as the number of species. Therefore, all species in this context can in principle coexist as a result of their interactions, without violating the competitive exclusion principle. Comparing communities with different ratio of positive versus negative influences in the initial pool, we also see that a community dominated with inhibitory influences has even lower chance of coexistence compared to the no-interaction control. As the ratio of facilitative versus inhibitory influences in the initial pool increases, the chance of coexistence also increases (Fig 2B, dotted versus solid versus dashed). We checked if members with the highest basal fitness have a higher chance of surviving: in communities with no interactions, that is indeed the case, but the pattern largely disappears in our interacting communities (Fig 2-FS2). Members with all basal fitness from the initial pool are represented in the final enriched communities (Fig 2-FS2), suggesting that interactions are a major factor in determining which species survive the enrichment process.

There are several assumptions in picking the initial pool of microbes. We asked how these assumptions might influence the coexistence outcome. The assumptions vary from large scale potential trends (e.g. the likelihood of facilitation versus inhibition) to detailed parameters (e.g. the rates of chemical release). Performing a complete search over the parameter space is impractical. Instead, we changed one parameter at a time and picked examples across the spectrum of possibilities to discover potential trends. For example, as we have shown in Fig 2B, the initial distribution of interaction types (facilitation versus inhibition) biases the outcome. Unfortunately, the available experimental data (e.g. (66, 68, 69) is not enough to definitively determine realistic assumptions for these assumptions. Examining interactions between microbes randomly sampled from the environment in a rich environment suggests that interactions are more likely to be inhibitory, rather than facilitative (70). However, since the experiments were performed in a rich environment with all necessary nutrients supplied, it is expected that competition for resources becomes dominant, underestimating cross-species benefit exchanges (71–73). Additionally, the likelihood of facilitative interactions in communities may be higher because of the role of facilitative interactions in community assembly (28). In the absence of a known general theme, we simulated initial pools with influences that are either mostly facilitative, or equally likely to be facilitative or inhibitory, or mostly inhibitory interactions. Our results show that positive influences encourages coexistence (Fig 2B).

We examined distributions of interaction strengths that were either less or more biased towards weak interactions in rich communities. Our results suggest that enrichment outcome is not sensitive to the details of the distribution of interaction strengths (Fig 2-FS3). Considering the difference between cases with and without interactions in Fig 2B, we asked how strong the interspecies interactions had to be to drive coexistence outcomes. Our results show that indeed when all interactions are chosen to be weak (relative to species’ basal fitness), coexistence are driven by neutral theory (Fig 2-FS4). We also observe that beyond some level, increasing the interaction strength does not further favor coexistence (Fig 2-FS4). This result suggests that when interactions are very strong, the coexistence outcome may be determined by the qualitative network structure (e.g. who facilitates whom), and not the quantitative interaction strengths.

To investigate how the network connectivity in the initial pool of species might influence coexistence, we examined networks with different architectures. For these simulations, we used a relatively strong interaction strength to ensure that interactions were the driving force of coexistence (Fig 2-FS4). First, using the binomial network architecture (in which each link having a fixed probability of being present), we investigated what would happen if nodes were more connected (i.e. more links per species and chemical). Note that this network architecture represents a situation in which all species on average are similarly generalists/specialists. We observe that in pools with many facilitative interactions, the coexistence outcome is insensitive to how many species each chemical influences (determined by probability ***q_c_***). More coexistence is achieved at intermediate levels of how many chemicals each species produces (determined by probability ***q_p_***) (Fig 2-FS5, left). In pools dominated by negative influences, lower connectivity favors coexistence as it reduces the chance of strong inhibitory influences (Fig 2-FS5, right).

What if the community consists of a mixture of specialist and generalist species? To answer this question, we changed the network architecture of the initial pool from binomial to power-law. In power-law networks, an architecture seen in many biological networks (74–76), there are few nodes that are highly connected (i.e. with many links), and many nodes with low connectedness (i.e. with few links). Our results show that power-law networks have the potential to produce higher richness in communities assembled through enrichment (Fig 2-FS6 and Fig 2-FS7). Specifically, we explored how coexistence with binomial versus power-law network architecture was affected when the number of species in the initial pool (Fig 2-FS6) or the number of mediators (Fig 2-FS7) were changed. Interestingly, the likelihood of coexistence is fairly insensitive to the number of species in the initial pool with a binomial interaction network, but pools with a power-law interaction network allow communities with higher richness, as the number of species in the initial pool increases (Fig 2-FS6). As the number of mediators in the initial pool increases, coexistence is fairly unaffected if the network is binomial, but power-law communities show less coexistence (Fig 2-FS7).

### 2.3. Depletable mediators enhance coexistence compared to reusable mediators

To address whether interaction mechanisms could impact coexistence outcomes, we examined communities with the same network of connectivity, but varied the fraction of species that consumed/degraded the chemical mediator (depletable mediator). We observe that if a higher fraction of interactions are through depletable mediators, coexistence becomes more likely (Fig 3).

**Fig 3.**
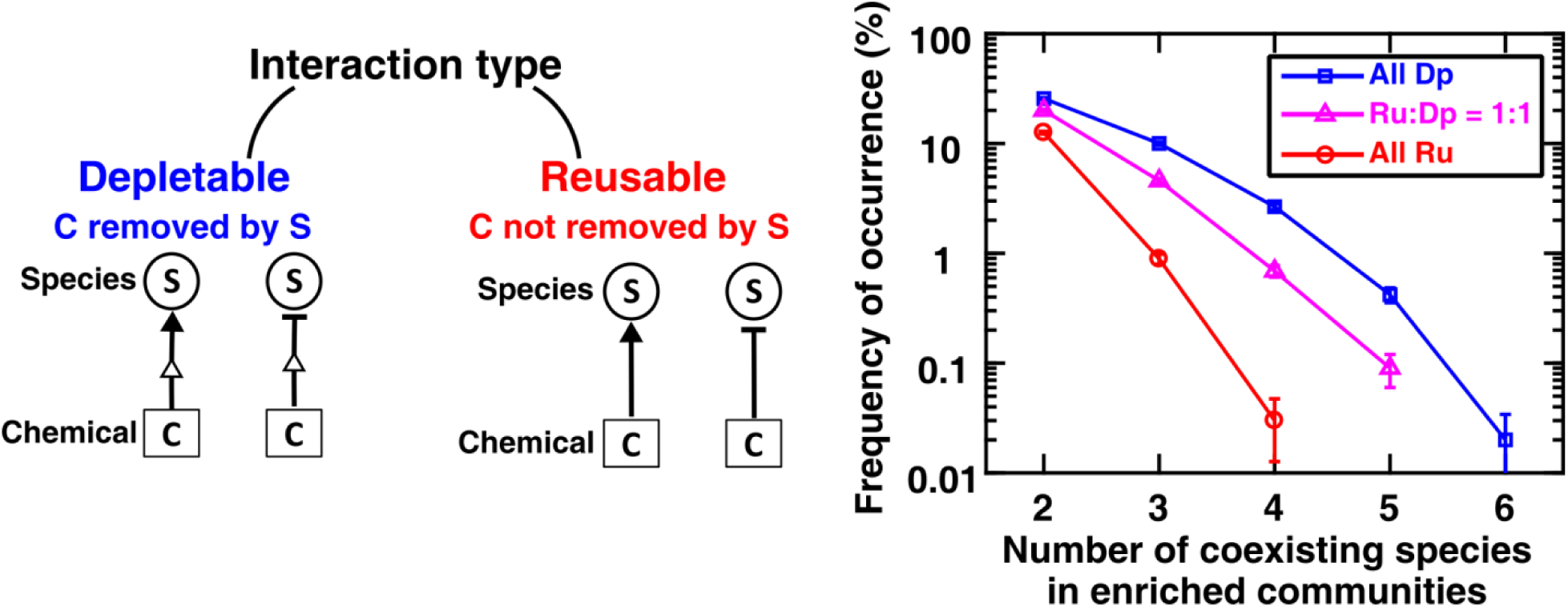
Interaction mechanisms impact coexistence. To explore how different interaction mechanisms might influence coexistence outcomes, we investigated two categories of chemical interactions: those with depletable mediators (Dp, in which recipients reduce the concentration of the chemical mediator by consumption or degradation) versus those with reusable mediators (Ru, in which recipients do not affect the mediator concentration). In the extreme cases, communities that used only Dp mediators showed significantly higher coexistence compared to communities that used only Ru mediators. We also examined enrichment in communities with equal number of the two interaction types, and coexistence in these “hybrid” communities appeared to be in between the two extremes. In these simulations, links had a binomial network. The initial number of species types N_c_=20, the number of possible mediators N_m_=15, and the ratio of interaction types +:-= 50%:50%.

This observed pattern of higher coexistence with depletable mediators seems to be consistent under a variety of conditions (e.g. Fig 2-FS6, Fig 2-FS7, and Fig 3-FS1). In Fig 3-FS1, we specifically examined quantitatively how coexistence was affected as we changed the ratio of the average rates of consumption to production of chemical mediators. With stronger consumption-to-release ratio (moving towards more depletable and less reusable mediators), more coexistence is achieved. This trend seems to saturate at very high ratios of consumption to production rates (Fig 3-FS1).

### 2.4. A pairwise model cannot accurately predict microbial coexistence

Since the difference between coexistence predictions in Fig 3 depended on interaction mechanisms, we asked whether a pairwise model, which by design does not include interaction mechanisms, would adequately predict coexistence of communities in which interactions are mediated through chemicals. Our previous studies on such communities (44) suggest that the standard Lotka-Volterra pairwise model is more likely to be suitable for communities in which interactions take place through reusable mediators. For this purpose, we establish an ecological network of pairwise interactions from interactions between each pair of species in the initial pool (44). We then simulate the enrichment process using both the “reference” model that incorporated the chemical mediators and using the corresponding pairwise model. Our results show that pairwise models fail to accurately predict coexistence in communities in which interactions take place through chemical mediators (Fig 4). Even though the overall patterns of richness likelihood may look similar and even though the pairwise model was fairly successful in predicting dynamics of each pair of species (see Methods for details), especially for reusable mediators, closer inspection shows that the coexisting species predicted by the pairwise model do not match the true coexisting ones from the reference model (Fig 4, bottom). The same overall observation is valid in communities derived from pools with more facilitative or inhibitory influences (Fig 4-FS1).

**Fig 4.**
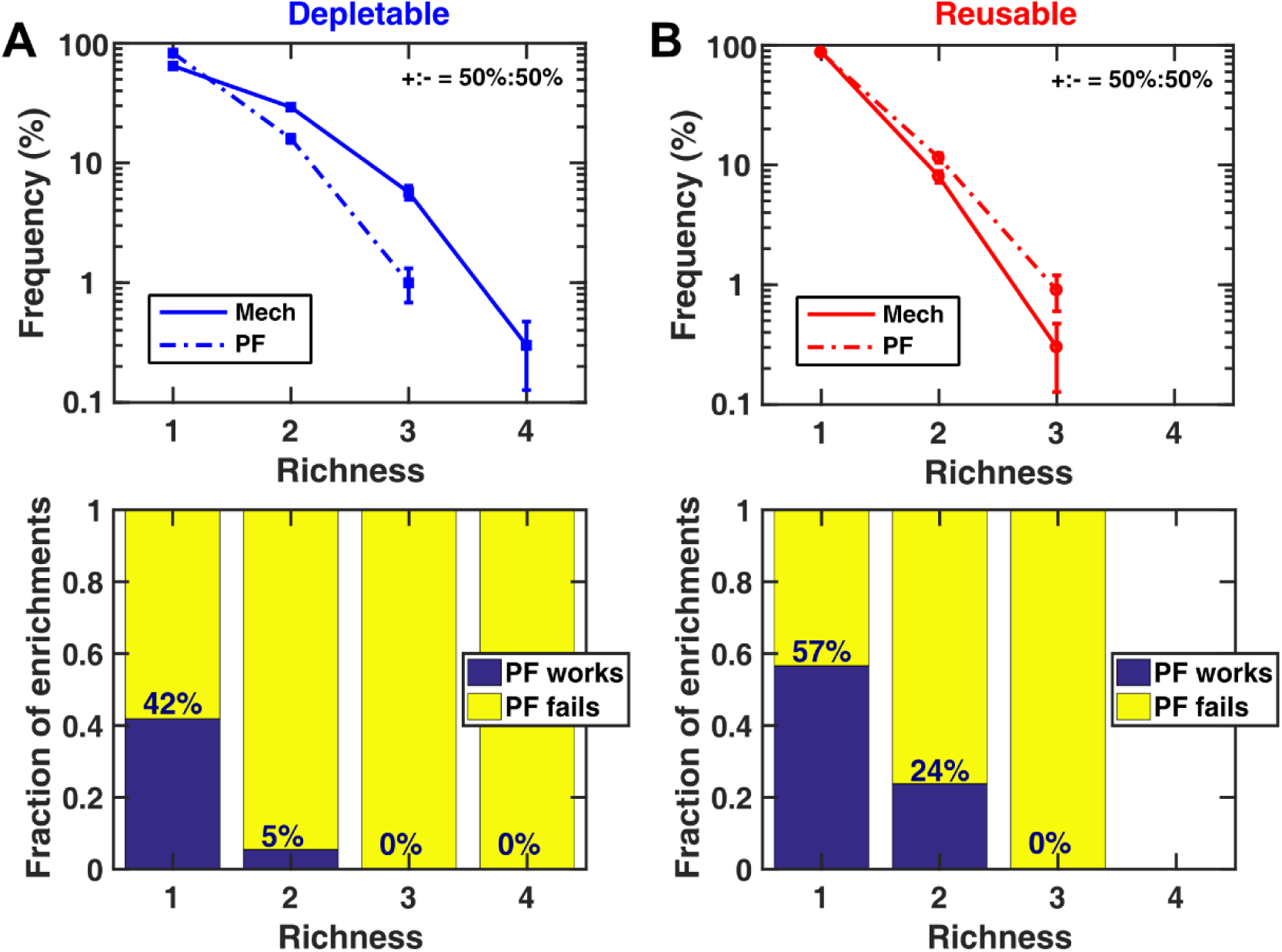
Pairwise models do not accurately predict coexistence. We compare richness likelihood obtained from the reference model that explicitly incorporates chemical mediators (Mechanistic, “Mech”) and that from a corresponding pairwise model derived for it (Pairwise Fitness, “PF”). The pairwise model is obtained based on characterizing interactions between each pair of species in the initial pool. We examined communities with only depletable mediators (A) or only reusable mediators (B). The pairwise model underestimates coexistence compared to the reference model in communities with depletable mediators and overestimates coexistence in communities with reusable mediators (top). Error bars show the sampling uncertainty. If we compare whether the pairwise model predicts the same species coexist as the reference mechanistic model, the pairwise model rarely succeed in producing correct predictions (bottom). In these simulations, the initial number of species types N_c_=7, the number of possible mediators N_m_=4, and the ratio of interaction types +:-= 50%:50%. The number of communities analyzed N_s_=1000. Other simulation parameters are listed in the Supplementary Information (Simulation parameters).

We speculate that the discrepancy between the predictions from a pairwise model and those of a more realistic model in Fig 4 stems from the shared environment and mediators. A pairwise model assumes that each interaction between a pair of species is independent of the presence of other members; however, that assumption would not be accurate when different community members produce and are potentially influenced by the same chemical mediators. We thus hypothesize that in a situation when such independence of interactions is more likely, a pairwise model would predict more accurate predictions. To test this hypothesis, we constructed examples with more number of chemical mediators, but fewer production and influence links per species. This will reduce the chance of cross-talk between different pairs of interacting species. Our results show that in such a situation, a pairwise model indeed provides more accurate predictions (Fig 4-FS2).

### 2.5. Facilitation and self-restraint are favored in enrichment

Among communities that showed coexistence, we searched for shared features. We categorized chemical influences based on whether they were facilitative versus inhibitory, and whether the species affected themselves versus other community members. Influences thus belong to one of four categories: self-facilitation, other-facilitation, self-restraint, and other-inhibition. We asked how enrichment for coexistence favored or disfavored each of these categories of influences. To answer this question, we compared how the frequency of each influence category changed from the initial pool to the final community of coexisting species. Our results suggest that in enriched communities, facilitation and self-restraint (i.e. production of chemicals that has an inhibitory effect on the producer) are favored (Fig 5A-B). Facilitation appears to be more prevalent in enriched communities, even if they are rare in the initial pool of interacting species (Fig 5B-C). This conclusion seems to be general and holds regardless of detailed parameters of the initial pool (e.g. when we vary the ratio of facilitative to inhibitory influences; Fig 5-FS1). Some enriched communities contained self-restraint and some did not (Fig 5C, left). Those without self-restraint all showed an increase in the frequency of facilitation during enrichment. Among those that contained self-restraint, almost all showed an increase in the frequency of self-restraint. The only communities that did not show a rise in frequency of facilitation still showed a rise in frequency of self-restraint (Fig 5). The same held true for other conditions explored (e.g. Fig 5-FS1).

**Fig 5.**
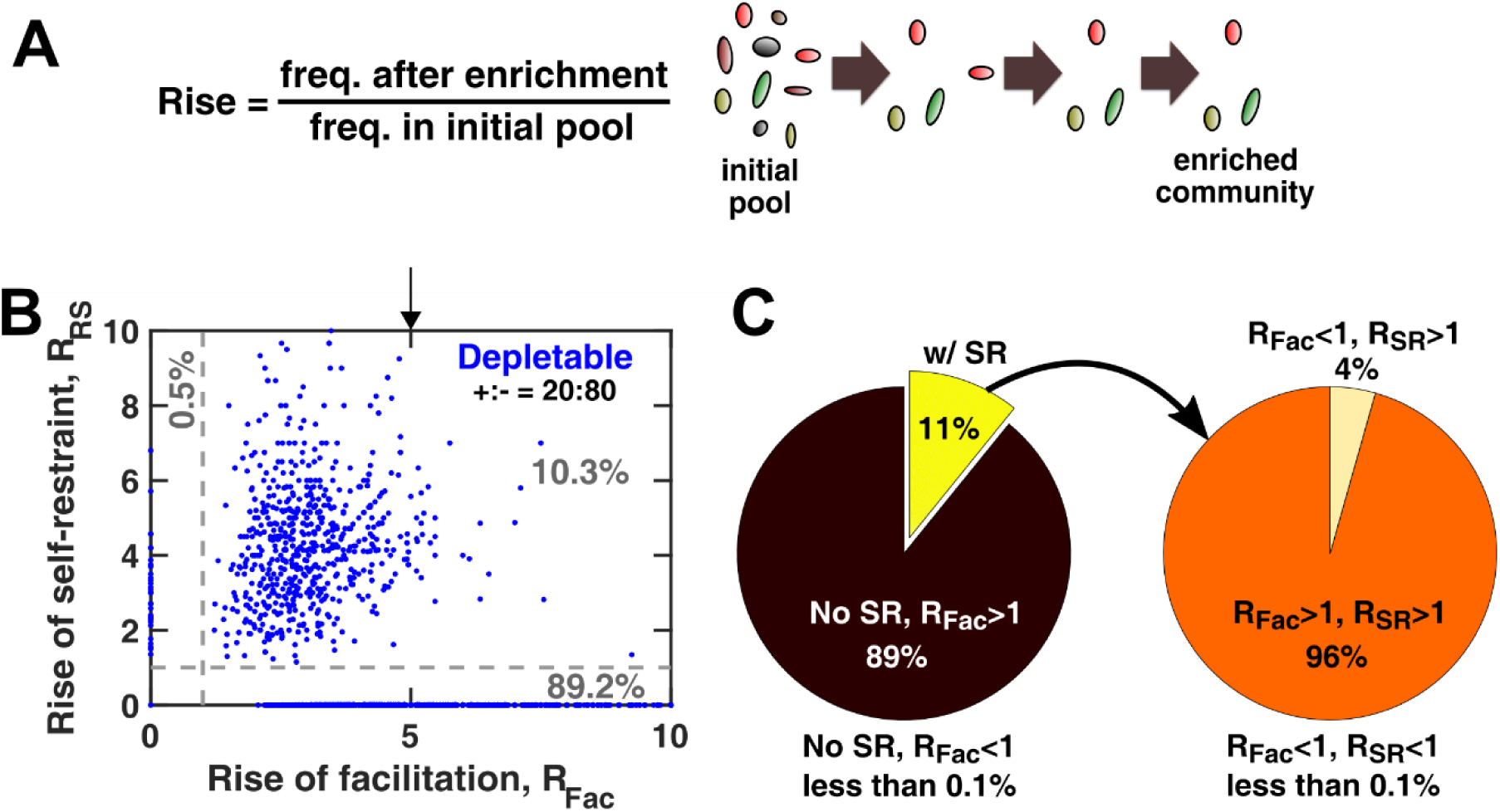
Facilitation and self-restraint are favored in enrichment. We define rise in frequency, R, as the change in the frequency of interaction types from the initial pool to the enriched coexistent community (A). Rise of frequency above 1 indicates that during enrichment, corresponding influences are more represented in final communities compared to the initial pool (see Methods). Examining the change in frequencies of facilitation and self-restraint reveals that these interactions are highly favored during enrichment (B). Most cases show an increase in the frequency of either facilitation or self-restraint (grey numbers show the fraction of data points in each region). We further examined the break-down of different categories in (C). Among all cases examined here, 11% of enriched communities contained self-restraint, whereas 89% did not. All of those lacking self-restraint showed an increase in the frequency of facilitation from the initial pool to the final community (R_Fac_>1). Among the 11% that contained self-restraint, all showed a rise in self-restraint above 1 (R_SR_>1), including 0.5% (4% within this case) that did not show a rise above 1 for facilitation. The number of communities analyzed N_s_=30000. In these simulations, the initial number of species types N_c_=20 and the number of possible mediators N_m_=15. Other simulation parameters are listed in the Supplementary Information (Simulation parameters).

The conclusion that facilitation and self-restraint arise as features of communities with coexistence is not surprising. The explanation for facilitation is intuitive: if a facilitative species rises in frequency, it improves the growth of its cohabitants and thus promotes coexistence. It is also intuitive that the negative self-feedback through self-restraint could prevent a species from outcompeting other members: as that species becomes more dominant, so becomes the inhibition it exerts on itself. This internal feedback, even if applied only to a few dominant members, can be the balancing force that allows coexistence.

Facilitation and self-restraint both have been suggested to play a role in coexistence and stability. From simpler two-species communities, we know that facilitation plays an important role in coexistence (25). Facilitation has also been implicated from field work on plant communities to increase community richness (77). Self-restraint is typically intrinsically assumed in pairwise models of ecological networks as a negative diagonal term in the matrix of interactions to incorporate the effect of intra-population competition (78). It is worth noting that in our analysis, the model was agnostic to this potential, yet self-restraint emerged as one of the features of enriched communities that exhibited coexistence.

### 2.6. Interspecies interactions causally impact coexistence

Considering that facilitation and self-restraint were correlated with coexistence, we asked if the relationship was causal. In other words, we would like to examine how important different influence categories are in coexistence during enrichment. To answer this question, we performed *in silico* “knock-out” experiments in which we remove an influence link from the community to examine how it affects final richness.

From the final enriched community, we picked one influence (chemical influence on a species) at a time, removed that influence from the initial pool of species, and asked how species richness was affected as a result. Our analysis shows that removing facilitative influences often leads to enriched communities with fewer species (Fig 6A). The same general pattern holds when we explore initial pools with other parameters. This shows that facilitation has a positive causal impact on coexistence. Among negative influences, self-restraint seems to contribute to coexistence more than other-inhibition, as removal of self-restraint more often leads to communities with lower richness (Fig 6A).

**Fig 6.**
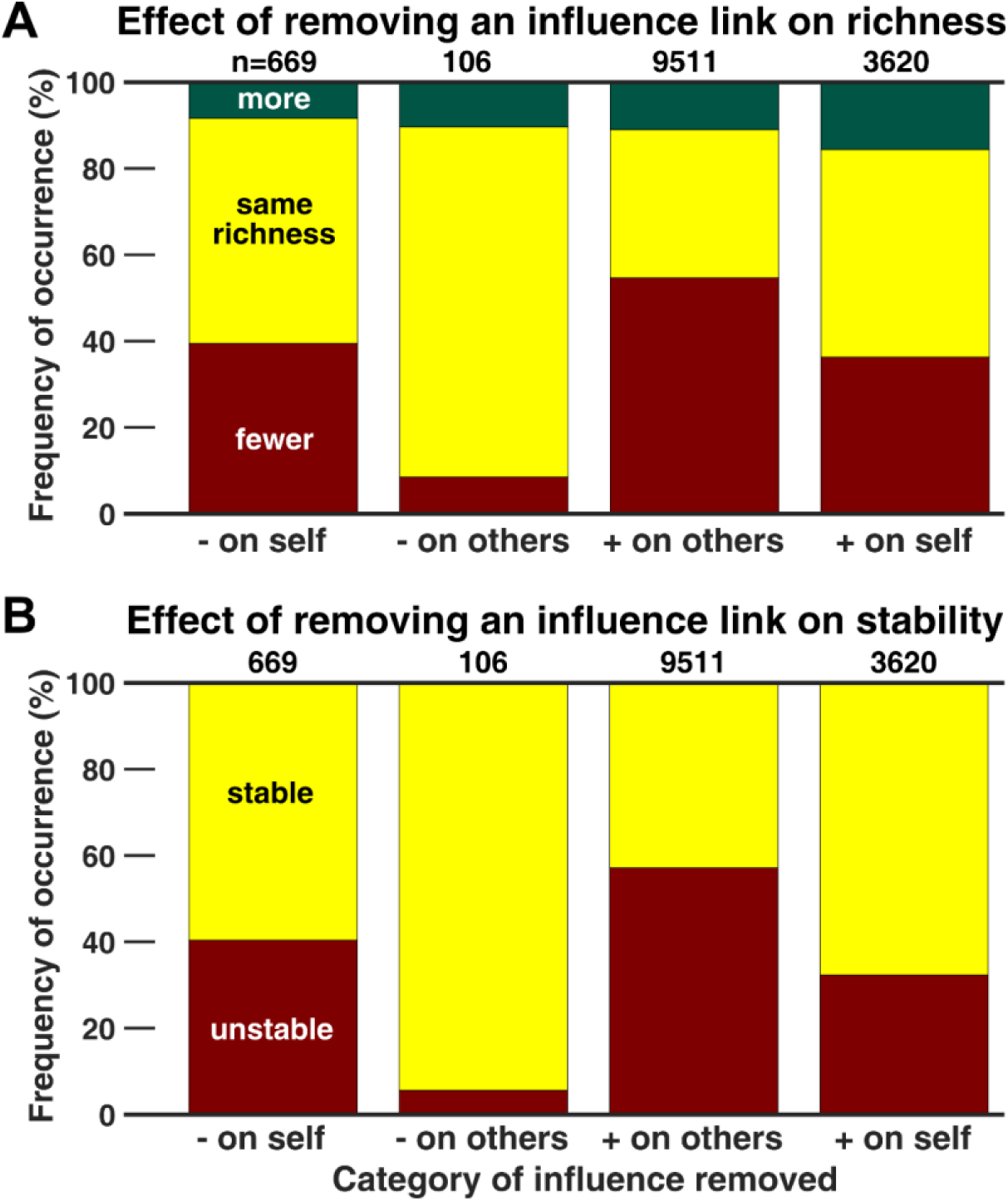
Even individual influences may causally impact coexistence and stability. To assess how interactions might influence coexistence, we use “knock-out” experiments in which we remove a link from the interaction network of a community that shows coexistence. In (A), the interaction is removed from the initial pool to ask how enrichment might be affected. We see that removing facilitation links has a considerable adverse effect on coexistence (with a large fraction of cases showing a drop in final richness). In contrast, removing self-restraint has a modest influence on coexistence, and removing inhibition of others on average benefits coexistence. In (B), the interaction was removed from the final community to see how an already stably coexistent community was affected by removal of different types of links. Observations in (B) match the trends in (A). Removal of facilitation likely disrupts the stable community, whereas removal of inhibition of others is unlikely to impact the community. Self-restraint interactions are at intermediate importance; their removal has a modest (~17%) chance of disrupting an already stable community in this example. Here, the initial pool has a binomial network and contains equally likely positive or negative influences (+:-= 50%:50%). Similar qualitative trends were valid when we examined communities with other sets of parameters.

We also started from the coexisting enriched community, removed an influence, and asked if the community remained stable afterwards. We functionally define stability by testing if we start from a community that has exhibited coexistence, after removing an influence, over the following 100 generations the same species coexist. We observed a trend similar to Fig 6A, in which removal of other-inhibition is least likely to disrupt stability, removal of self-restraint has intermediate chance of making the community unstable, whereas removing facilitation influences has a high chance of disrupting stability (Fig 6B).

Our interpretation is that during enrichment, some influences will remain in the community even though they are not necessary for coexistence. It appears that facilitative influences (and to an intermediate degree, self-restraint influences) are the necessary links that maintain the coexistence of species. In contrast, the remaining influences, especially other-inhibition ones that are disruptive to coexistence, “hitchhike” from the initial pool to the final enriched community. Removing these unnecessary influences is unlikely to make the community unstable.

## 3. Discussion

We refined a previous model of microbial interactions mediated by chemicals based on simple experimental investigations. We used this model in a computational screen to emulate the process of enrichment. By examining the commonalities among resulting communities that exhibited coexistence, we asked what features of their interaction might contribute to coexistence. We found that facilitation and self-restraint interactions played a critical role in allowing species coexistence. Importantly, this role was found to be causal in many cases: removal of a facilitation interaction negatively impacted coexistence, whereas, in contrast, more coexistence was likely observed after removing inhibition of other species. We also examined the effect of different parameters and revealed some of the factors that impact coexistence, including the interaction strength, the network architecture, and the role of depletable mediators. We found that our general conclusions (e.g. the role of facilitation in coexistence) were largely robust against changes to the details of parameters.

Our work re-emphasizes the importance of facilitation and self-restraint in community formation and maintenance (77, 79). Examples of how microbes employ these mechanisms for coexistence is not hard to imagine. Metabolic exchange has been considered as a common way other-facilitation can take place among microbes (80, 81), which would fit within the framework of our model. Self-facilitation can take place for example when a species breaks down complex compounds in the environment into consumable products. Practically this can take place by species that produce protease or cellulose; in our model, the mediator will be the product of the breakdown (e.g. amino acids or glucose). The concentration of such mediators increases in the presence of corresponding species, and those species (and potentially others) will experience a benefit in the presence of such products. Having unique access to these (otherwise inaccessible) resources allows these species to gain a fitness benefit that allows them to persist for coexistence. Self-restraint is also widespread, as the products of metabolism (such as acetate (82) or ethanol (21)) can often become inhibitory when they accumulate in the environment.

We also would like to emphasize that our model suggests that coexistence is affected by the mechanism of interactions among microbes (e.g. interactions mediated through depletable versus reusable chemical mediators). Our earlier work (44) had suggested that pairwise models that do not capture interaction mechanisms fail to properly capture community dynamics. That conclusion is re-iterated in this work where pairwise models failed to provide acceptable predictions of coexistence, except under very specific conditions in which all interactions were mediated by independent, reusable chemicals (consistent with the assumptions of pairwise models). We argue that for a community interacting through chemical mediators, pairwise modeling has two vulnerabilities: it ignores the mechanism of interaction (even though interactions can be through consumable versus reusable mediators) and it assumes that interactions between each pair of species are independent of other interactions (even though shared chemical mediators can be consumed, degraded, or produced by multiple species in a community). Our model of interactions through mediators addresses and resolves both these vulnerabilities. To be precise, our analysis does not conclusively assert that pairwise models are unfit for modeling coexistence. Nevertheless, the basic assumptions of our reference model are simple, intuitive, and commonly expected. The failure of pairwise models to make accurate predictions against our reference model (potentially representing a microbial community), should be considered a warning sign against the use of these models to predict microbial coexistence.

Do coexistence predictions from our model (i.e. the chance of achieving coexistence starting from a pool of microbes) match experimental observations? Different dependencies sampled in our exploration of the parameter space (Fig 2, Fig 2-FS4, Fig 2-FS5, Fig 2-FS6, Fig 2-FS7, Fig 3, and Fig 3-FS1) highlight the difficulty of experimentally validating the predictions. A rigorous experimental validation requires a well-characterized system with known interactions and chemical mediators. However, currently available experimental studies lack this level of detailed characterization. Examining some of the examples of reported coexistence starting from 2- and 3-species pools show fairly diverse outcomes (Fig 4-FS3). We performed experimental enrichment studies as well with different trio combinations of isolates from human nasal cavity, and observed no coexistence (data not shown). With our typical simulation parameters, we observed levels of coexistence within the range of previous experimental observations. Detailed evaluation of expected coexistence requires a more thorough investigation, which is beyond the scope of this work.

One of the intriguing observations in our results is the rapid drop in the likelihood of arriving at enriched communities with higher richness, especially with a binomial network architecture. Considering that in experimental settings instances of enriched communities with several species are not rare (63, 65, 83, 84), other factors besides fitness-changing interactions through chemicals may be in play. Some of these factors have been discussed before with potential impact on coexistence. Temporal or spatial heterogeneity (20, 85) could be a factor that effectively changes the strength of interactions among community members over time. Additionally, shared resources are assumed to be always in abundance in our model; however, if some resources become scarce, species with lower intrinsic fitness might be favored, leading to increased coexistence.

In modeling microbial coexistence, there are many other features that deserve a closer look. Species behavior and physiology may change depending on the environmental cues or intercellular communications (86)(Hart et al 2018). There may be an intrinsic heterogeneity, in terms of microbial phenotypes, within each population (87). Another possibility is potential intrinsic structure in the network of the community (e.g. presence of intrinsic modularity), which may cause the coexistence to deviate from predictions of our model based on randomly assigned networks. Lastly, our ecological model does not incorporate evolutionary changes in species, which could potentially impact coexistence outcomes. These factors are outside the scope of this report, but can be independently examined using the same framework in the future. Other factors such as non-monotonic interactions (88), and non-additive interactions (46) are some of the aspects that can influence the formation and maintenance of microbial communities, but are beyond the scope of our work. We hope that this work will be a stepping stone in formulating and capturing important features of microbial interactions in community models.

## Methods

### Simulation platform

Simulations were implemented in Matlab^®^ and run on the shared Scientific Computing cluster at the Fred Hutchinson Cancer Research Center. The cluster is a Linux-based 64-bit computing platform with 64GB of RAM per job. Typically, different sets of assumptions and conditions were screened and then analyzed to find trends in networks of enriched communities. Simulation codes are available in the supplementary files associated with this manuscript.

### Network architecture of initial species pool

In binomial networks (Fig 2-FS6 and Fig 2-FS7), the presence or absence of c-links and f-links each is determined by a fixed probability. In power-law networks (Fig 2-FS6 and Fig 2-FS7), the number of c-links per species and the number of f-links per chemical follow a power-law distribution. The basal fitness values of species in the initial pool of species is picked randomly from a uniform distribution (r_0_ ~ U(0.08,0.12) per hour). The exact value of basal fitness is inconsequential as all other fitness values (e.g. fitness influence of interactions) and time-scales (e.g. update time-step) can be scaled accordingly without loss of generality. For interaction strengths, the values within the main manuscript are picked randomly from a uniform distribution when the fraction of positive to negative interactions is 1:1. In cases where either positive or negative interactions are more likely, the absolute values of interaction strengths within positive or negative interactions still follow a uniform distribution, but the sign will be positive or negative based on a binomial distribution.

### Experimental characterization of chemical facilitation and inhibition

For facilitation, we examined growth of *E. coli* K12 MG1655 single gene knockout auxotrophic strains in media supplemented with the corresponding amino acid at different concentrations. For leucine auxotrophy, we replaced LeuB with a chloramphenicol resistance gene and for isoleucine auxotrophy, we replaced IleA with a kanamycin resistance gene. For isoleucine auxotrophs, a BioTek Synergy Mx multi-mode microplate reader was used to monitor the optical density (OD) cells over 24 hours at 5 minute intervals. Cultures typically started from an initial OD of 0.001, and were kept shaking in between OD readings. Standard M9 minimal medium (89) was used as the basal growth medium in these experiments, and it was supplemented with isoleucine as needed. For leucine auxotrophs, the OD assay above was not sensitive enough to measure the growth rate. Instead, we used a fluorescently labeled strain (using DsRed on a plasmid) and used the plate reader to monitor the total fluorescence from the cultures growing when supplemented with different concentrations of leucine. Excitation was set at 560 nm and emission at 588 nm in this assay. We used only the first 3 hours of the fluorescence reading to calculate the growth rates to minimize the effect of leucine depletion as cells were growing.

For inhibition, we examined different combinations of bacterial strains and inhibiting compounds, as listed in the following table:

| Strain | Inhibitor | Medium | Temperature |
| --- | --- | --- | --- |
| *E. coli* K12 MG1655 | Acetic acid | M9 | 37°C |
| *E. coli* K12 MG1655 | Erythromycin | M9 | 37°C |
| *Brevibacillus* M1-5 | Acetic acid | BAAD | 50°C |
| *Staphylococcus aureus* SD6 | Acetic acid | 10% THY | 37°C |
| *Staphylococcus aureus* SD6 | Erythromycin | 10% THY | 37°C |
| *Staphylococcus epidermidis* SD8 | Acetic acid | 10% THY | 37°C |

For these inhibition experiments, we typically streaked them on rich medium (on LB for *E. coli* strains, on PCS for the *Brevibacillus* strain, and on BHI for *Staphylococcus* strains) and isolated a clone. The clone was then grown to exponential phase in the basal media listed in the table, in the absence of inhibitors. Multiple replicate wells on either a 96-well plate or a 384-well plate were inoculated with these exponentially growing cells typically at an initial OD of 0.0012, at different concentrations of the corresponding inhibitor. Growth was monitored by recording the OD at 5-min or 10-min intervals using either a BioTek Synergy Mx multi-mode microplate reader, or a BioTek Epoch2 absorbance microplate reader. Plates were incubated while shaking inside the plate reader in between OD readings. Typically 3-6 replicates were used per condition. The wells around the periphery of microplates were found to be more subject to evaporation. We thus filled those with sterile water to reduce the impact of evaporation on other wells and only used the rest of the wells on each plate for our cultures.

### Analyzing the growth rate from experimental OD readings

To estimate what the growth rate is in each well, we exported the data from Gen5 software that controls microplate readers to a text file, and transferred the data to Matlab for analysis. For each well, we used the wells in time-points 3-10 to estimate the background OD corresponding to that well. The first two time-points were dropped, because we occasionally saw condensation issues before the plate reached the incubation temperature. After subtracting the background, we picked data points for each growth curve that were between OD values of 0.002 and 0.02 to avoid noise at low ODs and saturation at high ODs. A linear function was then fit into the log of OD values using the polyfit function in Matlab. The slope of this line was reported as the growth rate for that well.

### Fitting a pairwise model into the dynamics of two species growing together

To model enrichment using a pairwise model, we build a network by characterizing the interaction between each pair of species. For each pair of species, we first simulate monocultures using the mechanistic model and estimate the parameters that approximate the growth of each monoculture (using DeriveBasalFitnessPastTransient_WM_DpMM.m). To derive a pairwise model for cocultures of two species, we start from an initial ratio of 1:1 and simulate the dynamics for 30 generations. We use the last 10 generations of this simulation as the “fitting window” to estimate the pairwise model parameters (using DeriveSatLVPastTransient2_WM_DpMM.m). This allows us to exclude the transient effects known to interfere with parameter estimation (44). If the ratio of the two species at the beginning of this window drop below 1:1000, we adjust the initial ratio to counter that. This simulates the experimental process of adjusting population ratios meant to allow both populations to be present at a large enough ratio for reliable estimation of their influence on each other. Given 10 generations of population growth in which both species are represented, we use the nonlinear least square optimization routine (lsqnonlin in Matlab®) to find four parameters of a canonical pairwise model that offer a best fit for population dynamics. The canonical pairwise model (44) used here can be described as

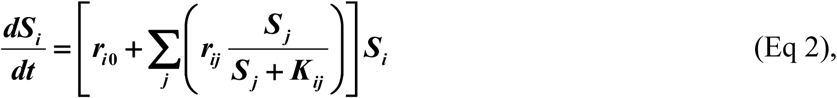

in which *r_i0_* is the basal fitness of species *i*, *r_ij_* is the maximum fitness effect exerted on species *i* by species *j* in their interaction, and the interaction follows a saturating form ***S _j_/(S_j_+K_ij_*)**. We use the residual error output of this optimization routine to assess whether the fitting is successful or not (with typical parameters, residual < 200 is assumed to be a successful fit).

### Calculating the rise in frequency of different interaction types

We define the rise in frequency as the change in the frequency of interaction types from the initial pool to the enriched coexistent community. In our calculations, we screened the enriched communities to ensure their network had a connected graph, meaning that all species either were influenced by a chemical produced by other species or produced a chemical that influenced other species. This screening was performed to remove communities that achieved coexistence without interactions. Additionally, we removed all interactions with a fitness influence lower than 1/500^th^ of the average basal fitness. The motivation was that such interactions would be inconsequential within the 300-generation time-scale of our enrichment experiment. Removing these spurious links in the graph would thus make it easier to see the patterns for more important interactions.

## Supplementary information

List of Codes: Matlab scripts used for modeling and analysis of data are listed below

| Code | Description |
| --- | --- |
| EnrichmentComparison_WM_Mech_vs_PF2_ABE_RD.m | Compares enrichment predictions between the mechanistic model (chemical mediators explicitly incorporated) versus an estimated pairwise model derived from it. |
| Enrichment_WM_DpMM.m | Simulates the enrichment process for communities with given parameters using the model that explicitly includes chemical mediators. |
| Enrichment_WM_PF.m | Simulates the enrichment process for communities with given parameters using a pairwise model. |
| DeriveBasalFitnessPastTransient_WM_DpMM.m | Uses an optimization routine to estimate model parameters that provide a best fit into growth dynamics of each species growing in a monoculture. |
| DeriveSatLVPastTransient2_WM_DpMM.m | Uses an optimization routine to estimate model parameters that provide a best fit canonical pairwise model into growth dynamics of two species growing in a coculture. |
| DistInteractionStrengthMT_PB.m | Produces a matrix of interaction strengths with uniform distribution of strengths and designated probability of positive versus negative signs. |
| Dynamics_WM_DpMM.m | Simulates the dynamics of a well-mixed community of species interacting via chemical mediators |
| Dynamics_WM_SatLV.m | Simulates the dynamics of a well-mixed community of species engaged in pairwise fitness interactions (saturable Lotka-Volterra type (Eq. 2)) |
| FitCost_BasalFitness.m | Cost function for fitting a saturable LV model (Eq. 2) into a given population dynamics of a single species. |
| FitCostGR2_SatLV.m | Cost function for fitting a saturable LV model (Eq. 2) into a given population dynamics of two species. |
| NetworkConfig_Binomial.m | Produces matrix of links for a network with binomial connectivity |
| NetworkConfig_Powerlaw.m | Produces matrix of links for a network with power-law connectivity |

**Fig 1-FS1.**
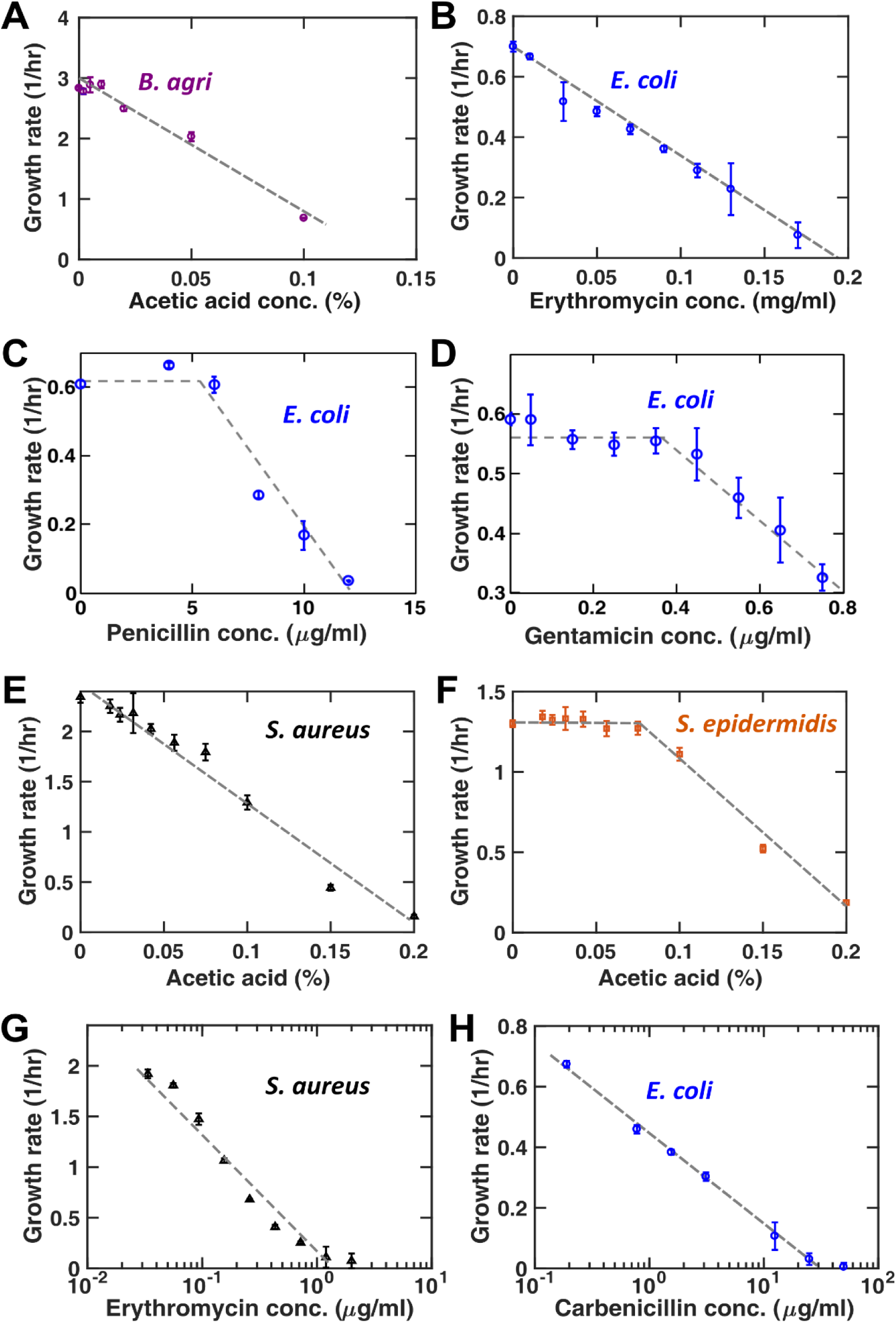
Inhibition by chemical compounds often shows a consistent pattern of linear decrease, beyond a threshold concentration. Examining the growth inhibition by chemical compounds in many cases we have examined showed a linear decrease as the inhibitor concentration increased. (A) and (B) show two examples of this trend for *B. agri* and *E. coli*, in response to acetic acid and erythromycin, respectively. In these cases we observed a small threshold effect (i.e. no inhibition or slight growth rate increase at very small concentrations of the inhibitor), however, the threshold was low enough that a linear decrease in growth rate, i.e. ***dS/dt =(r_0_−ηC)S*** offered a good approximation. (C) and (D) In some cases, we observed a more pronounced threshold effect, for example when *E. coli* was exposed to penicillin or gentamicin. This means these bacteria can tolerate the inhibitor to some extent, but beyond the threshold concentration, ***Cth***, cells will be affected by the chemical inhibitor. For these situations, a more accurate model is 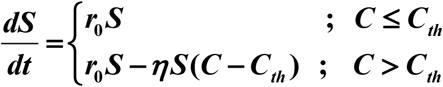. (E) and (F) Other species and strains showed the same overall trends in many cases. As representative examples, growth rates of *S. aureus* and *S. epidermidis* in response to acetic acid show linear decrease (similar to (A-B)) and linear decrease beyond a threshold (similar to (C-D)), respectively. (G) and (H) Although the trends in (A-F) were most common, a third pattern was observed at lower frequency in which the decrease in growth rate was proportional to the logarithm of the inhibitor concentration. For simplicity, we chose the more common linear decrease trend in our simulations.

**Fig 1-FS2.**
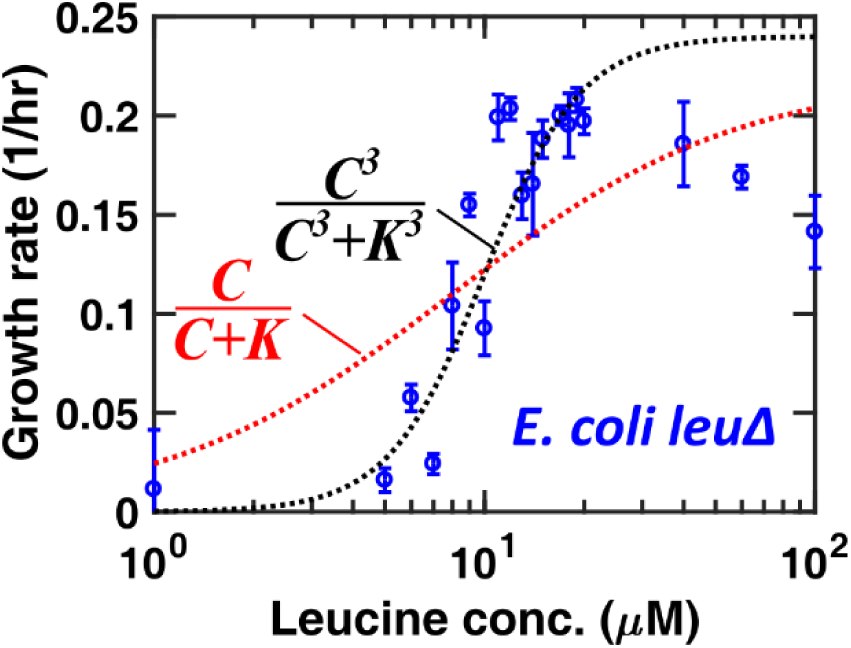
Facilitation by chemical compounds shows a consistent pattern of saturable response. Similar to Fig 1D, we have observed that the influence of a facilitative chemical compound on species can be approximated by a Moser growth model. In this case, a leucine auxotrophic strain of *E. coli* is grown in defined M9 media supplemented with different concentrations of leucine. To avoid potential effects of leucine depletion, growth in the first 3 hours of growth is used to estimate the growth rate. Experimental observations of this auxotrophic strain suggest that a third-order relation (black dotted line) offers a more accurate estimation compared to the first-order Monod-type equation (red dotted curve). For simplicity, the first-order Mono-type equation is used in the model which still captures the saturating form of the equation. Similar to Fig 1D, at much higher concentrations of leucine, potential toxic effects reduce the growth rate (above 50 μM of leucine), but we do not include this effect in our model.

**Fig 2-FS1.**
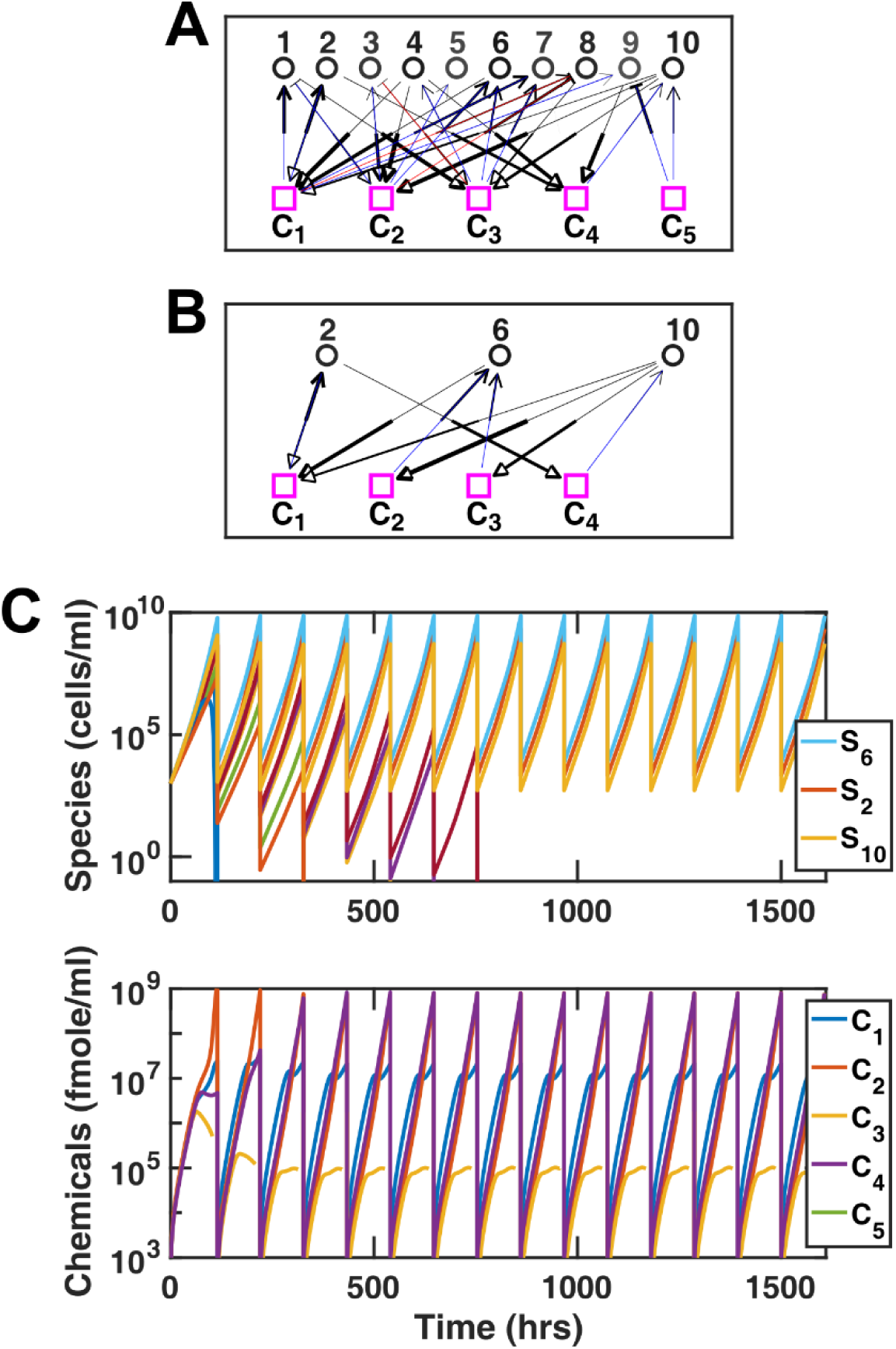
During enrichment a subset of species can achieve coexistence. We show an example of the dynamics and network of a community assembled through enrichment. In (A) the random network of interactions in the initial pool of 10 species and 5 mediators is shown. (B) After enrichment, 3 species and 4 mediators remain in the community. In (A) and (B), the thickness of f-links indicates their relative strength and the thickness of c-links indicates the relative rate of removal or production. The shades of species indicates their relative basal fitness (darker shades for higher basal fitness values). Additionally, for f-links, those removed by their recipients are marked as blue (depletable) and those not removed by their recipients are marked as red (reusable). In this example, the initial ratio of positive to negative influences is 80%:20%, and the ratio of depletable to reusable mediator links is 80%:20%. (C) Dynamics of species population sizes (top) and chemical concentrations (bottom) are shown for the example of transition from (A) to (B). All populations are assumed to have a similar size at the beginning of enrichment. Any population that drops below 0.1 cells/ml is assumed to be extinct and is removed from the rest of the simulation. In this example, 3 species achieve coexistence, and coexistence appears to be stable after 7 rounds of growth-dilution.

**Fig 2-FS2.**
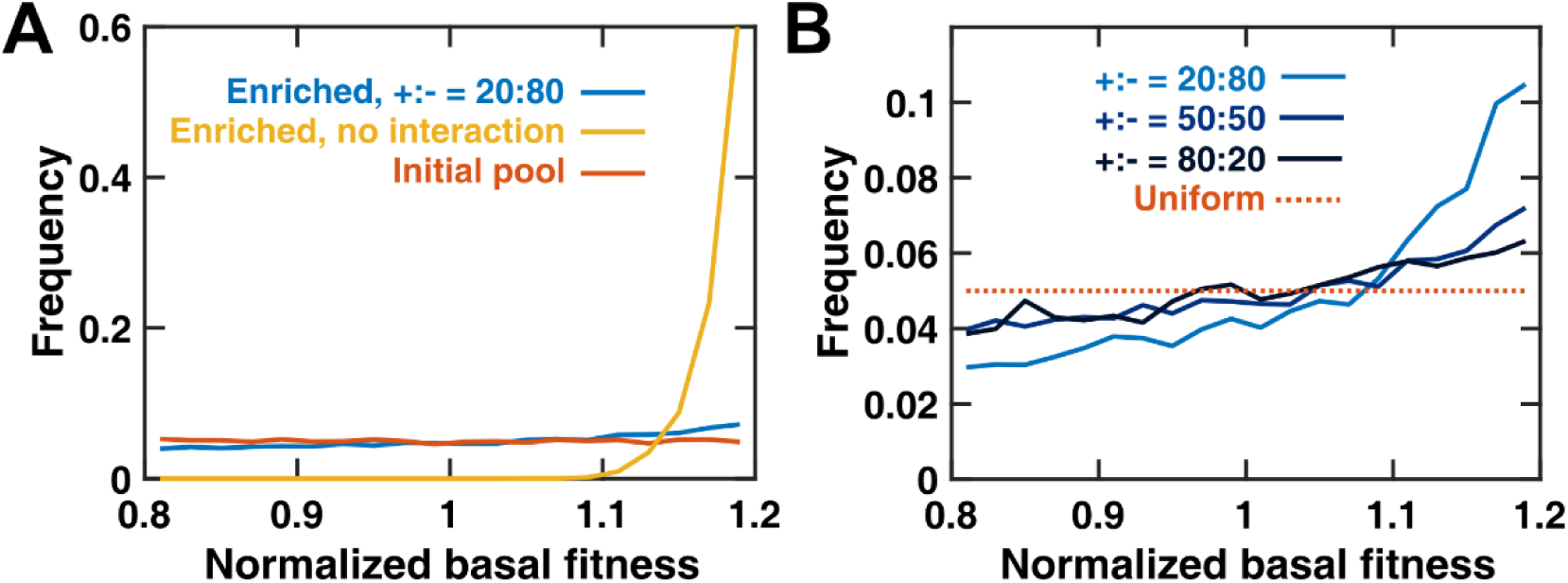
Basal fitness is a major determinant of coexistence in the absence of interactions, but not when species are engaged in strong interactions. (A) Examining the distribution of basal fitness in all enriched communities, we observe a slight advantage for species with higher basal fitness in communities obtained from a pool of interacting species. In the absence of interactions (representing neutral theory), species with the highest basal fitness will outcompete other species. The distribution in the initial pool is uniform across all basal fitness values. These distributions are plotted as histograms of frequencies over 20 bins. (B) Comparing the distribution of basal fitness in enriched communities, we observe that although higher basal fitness increases the chance of being present in enriched communities, this trend is more pronounced if initial pools were dominated by inhibitory influences compared to facilitative influences. The dotted line shows the theoretical distribution of basal fitness values in the initial pool. These distributions are plotted as histograms of frequencies over 20 bins. In simulations in (A) and (B), the average interaction strength relative to average basal fitness, r_i_/r_0_ = 2. The number of coexistent cases examined is 5000.

**Fig 2-FS3.**
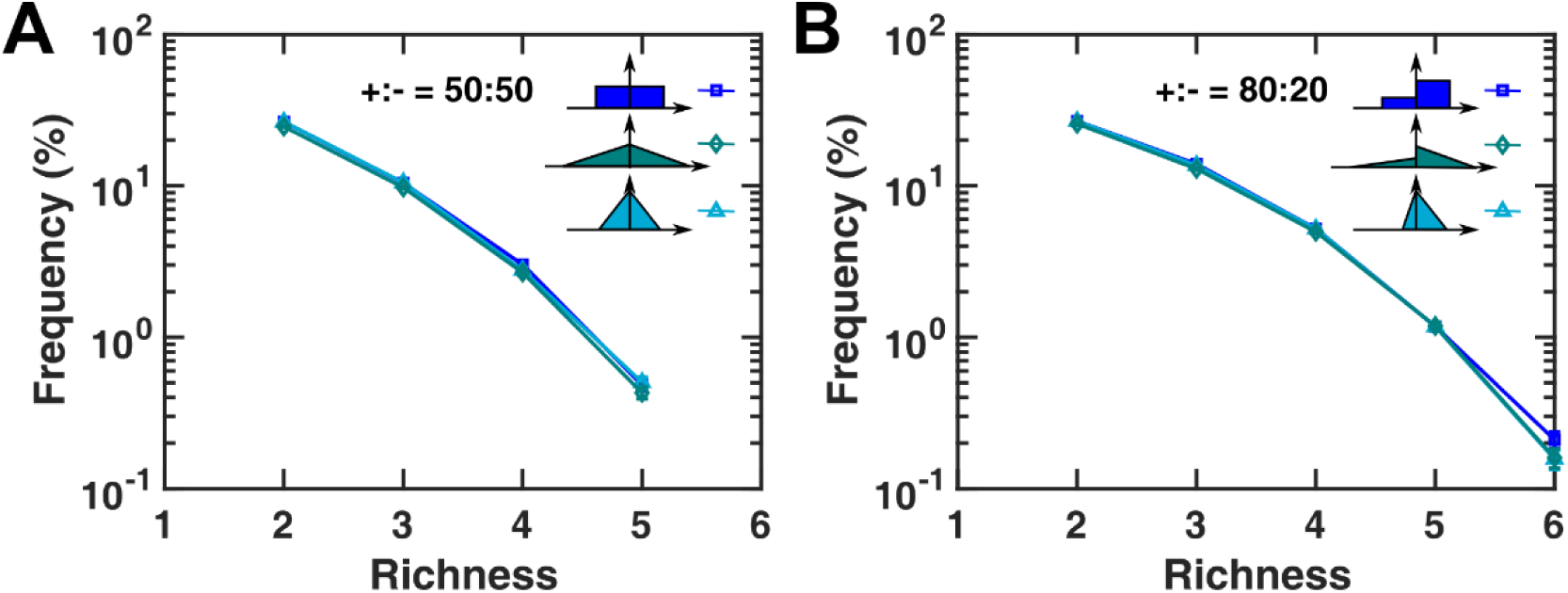
Coexistence outcome is not sensitive to the distribution of interaction strength values. Examining how the distribution of interaction strengths in the initial pool affects coexistence, we find that the likelihood of coexistence is only modestly affected by the distribution of interaction strengths. We examined situations where strong and weak interactions were equally likely (blue squares), where strong interactions were less likely (green diamonds), as well as a situation in which the less likely interaction type (positive or negative) has lower maximum interaction strength (cyan triangles). Comparing the frequency of arriving at communities with different richness through enrichment shows a negligible difference in the outcome when different interaction strength distributions are used when assigning interactions in the initial pool. Only simulation results with depletable mediators are shown, when (A) positive and negative influences are equally likely, and (B) positive influences are more likely than negative influences. In these simulations, the average interaction strength relative to average basal fitness, r_i_/r_0_ = 1.

**Fig 2-FS4.**
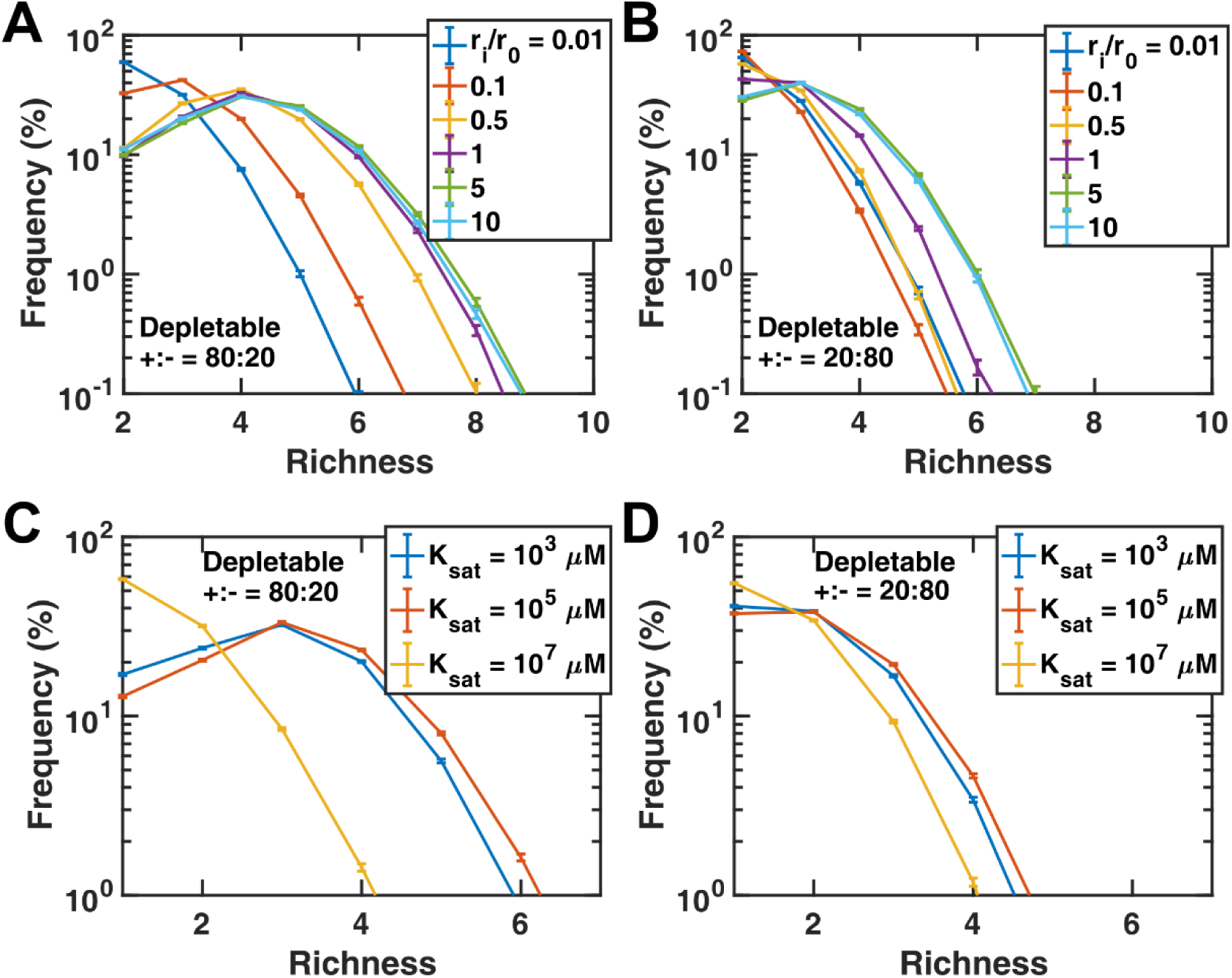
Coexistence is disrupted at weak interaction strengths. We examined how interaction strength affects coexistence. Whether influences are mostly facilitative (A) or inhibitive (B), as interactions become stronger (i.e. larger r_i_/r_0_ values, where r_i_ is the average interaction strength magnitude and r_0_ is the average net basal growth rate), coexistence becomes more likely, but this trend saturates at very strong interactions. At very weak interaction strengths, we can assume that coexistence happens only for species with very similar growth rates, approaching what is expected from the neutral theory. (C-D) As we vary K_sat_, the average of saturation concentration K_i,l_ values, we observe a trend similar to varying the interaction strengths. Large values of K_sat_ effectively represent weaker interactions (see Eq. 1) and lead to lower likelihood of interaction-driven coexistence, as expected.

**Fig 2-FS5.**
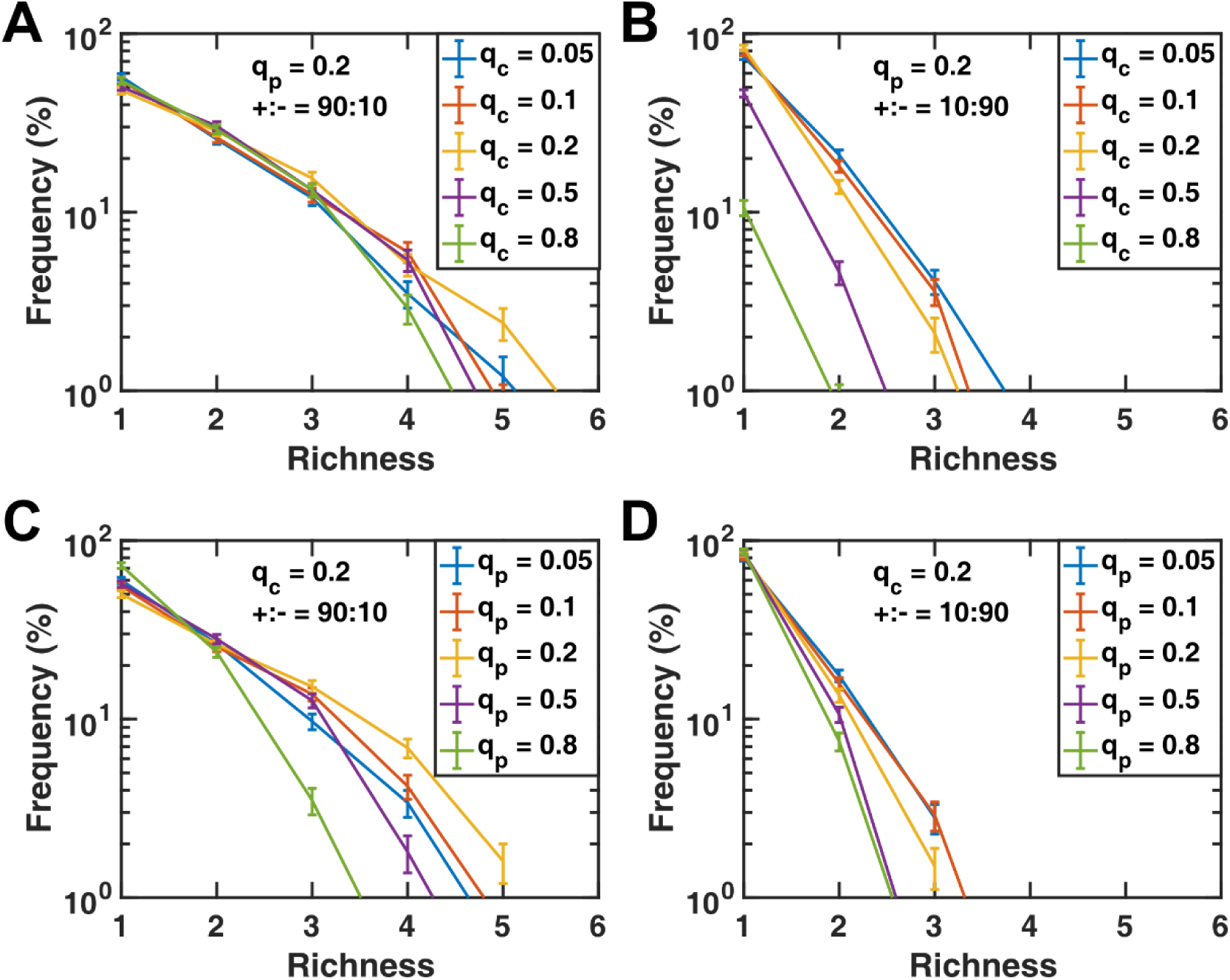
The degree of connectivity per node can impact coexistence. We examined how the degree of connectivity of species and chemicals influenced coexistence. For this purpose, in a binomial network, we (A-B) changed the chance of presence of influence links that affect species fitness **(*q_c_*)** and (C-D) the chance of presence of production links **(*q_p_*)**. Our results show that when species are engaged mostly in facilitative interactions, coexistence is favored at intermediate levels of connectivity (A and C). In contrast, in communities dominated by inhibitory interactions, coexistence is favored at lower connectivity (B and D).

**Fig 2-FS6.**
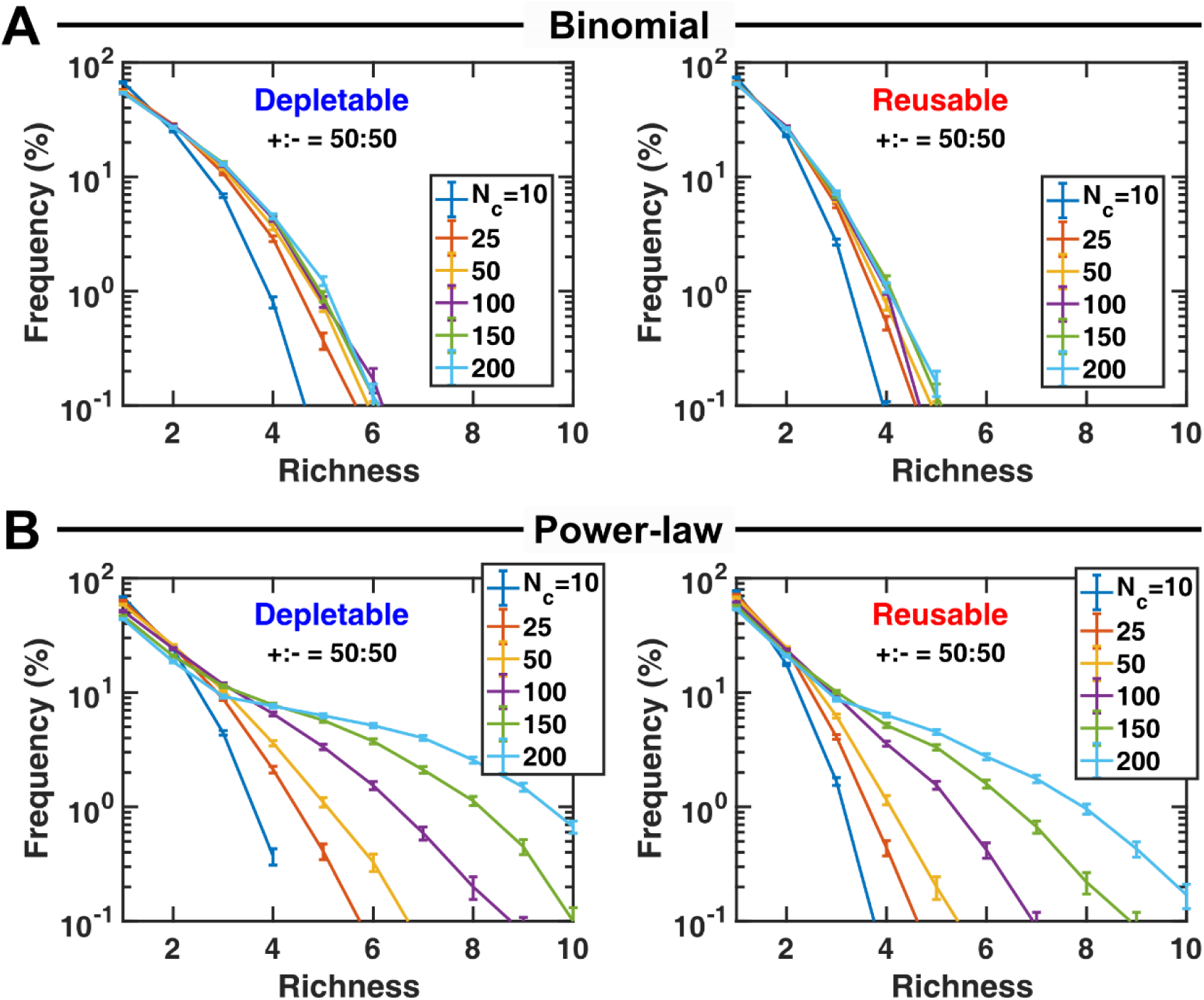
Network architecture impacts coexistence. We examined coexistence in communities with two different network architectures starting from an initial pool with different number of species. (A) In binomial networks (used in the rest of the examples in this paper), each chemical production link or species influence link exists with a fixed probability. (B) In power-law networks, the chance of being connected (each species to chemicals or each chemical to species) exponentially drops as the number of connections per node increases. As a result, most nodes have few connections, and a small fraction of nodes are highly connected. We observe that for power-law networks, it is more likely to obtain communities with multiple species, compared to binomial networks. In both cases, we observe more coexistence with depletable mediators, consistent with Fig 3. In these simulations, the number of possible mediators N_m_=15. For binomial networks, ***q_p_***=0.2 and ***q_c_***=0.2. For power-law networks, we choose the parameters such that the average connectivity is similar to our binomial networks. The number of communities analyzed N_s_=10000.

**Fig 2-FS7.**
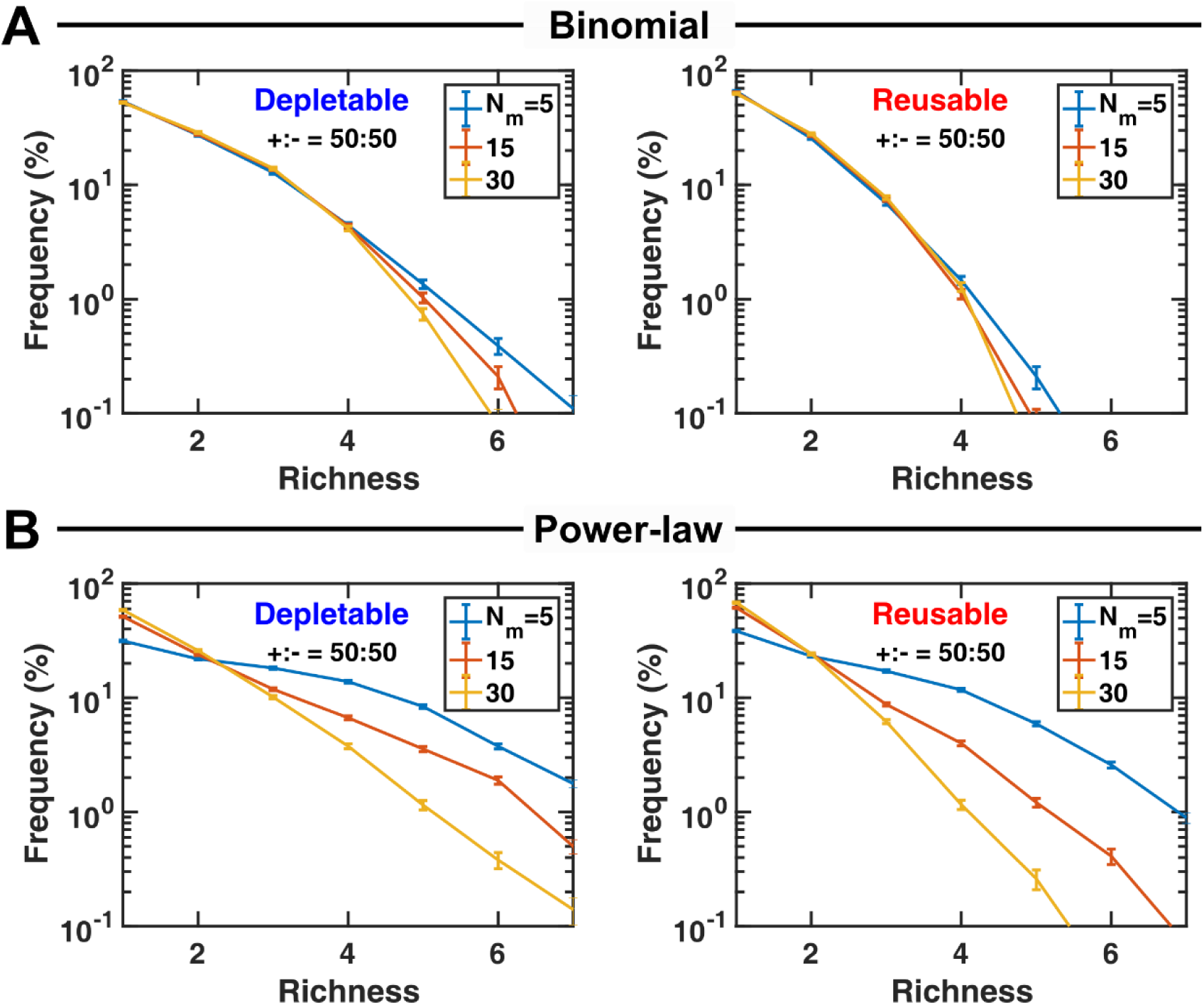
A power-law network architecture favors coexistence compared to a binomial network architecture. Similar to Fig 2-FS6, we examined coexistence of communities with different network architectures. We looked into how different number of mediators affected coexistence. We observe that binomial networks (A) are less likely to lead to coexistence compared to power-law networks (B). We also observe that with fewer mediators, the chance of coexistence increases, and this trend is more pronounced in communities with a power-law network architecture. Additionally, in both cases, we observe higher coexistence with depletable mediators, consistent with Fig 3. In these simulations, the number of species in the initial pool N_c_=100. For binomial networks, ***q_p_***=0.2 and ***q_c_***=0.2. For power-law networks, we choose the parameters such that the average connectivity is similar to our binomial networks. The number of communities analyzed N_s_=10000.

**Fig 3-FS1.**
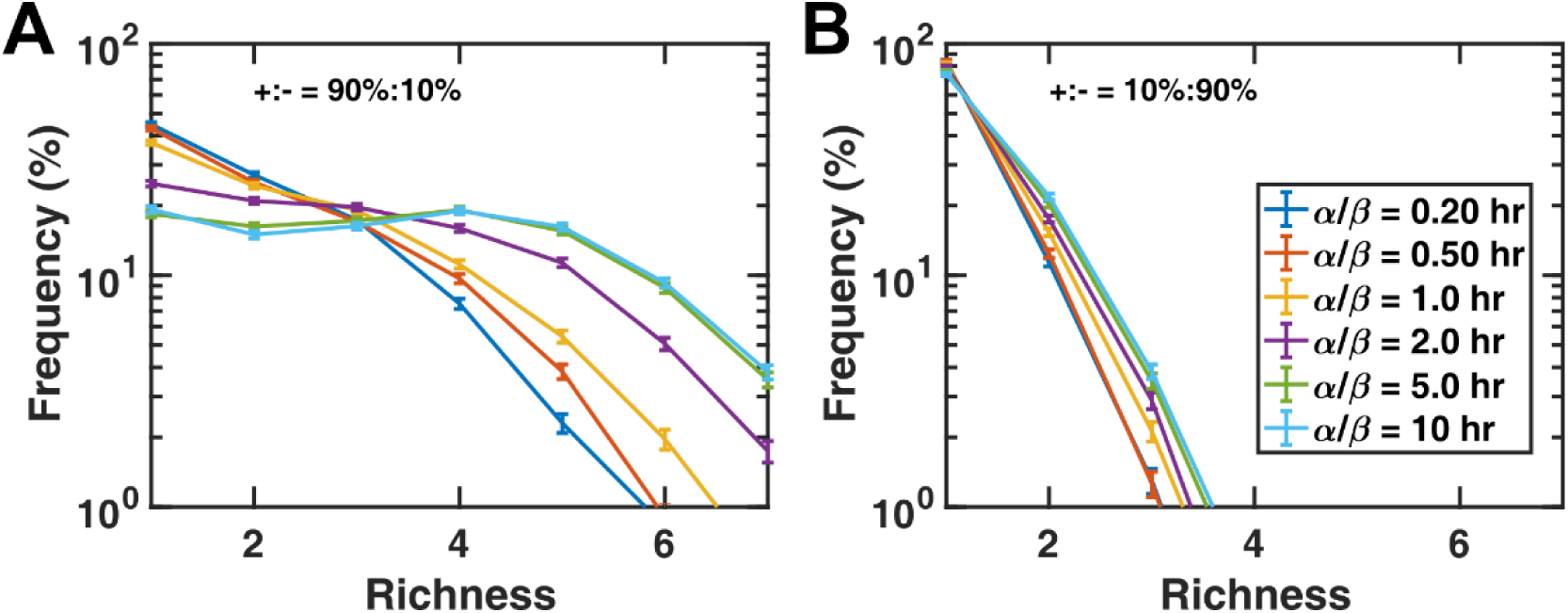
Depletion of mediators favors coexistence. We examined how changes in average consumption and production rates of chemical mediators (***α*** and ***β***, respectively) affected coexistence. For both communities that contained many positive influences (A) and those that contained many negative influences (B), an increase in the ratio of *α*/*β* increased the likelihood of coexistence. This trend was more pronounced in communities that had more positive influences (A), but was saturated beyond a ratio of 5. In these simulations, the initial number of species types N_c_=20, the number of possible mediators N_m_=15, and a binomial network connectivity is assumed with ***q_p_***=0.2 and ***q_c_***=0.2. The average basal fitness in the initial pool is 0.1/hr. The number of communities analyzed N_s_=5000.

**Fig 4-FS1.**
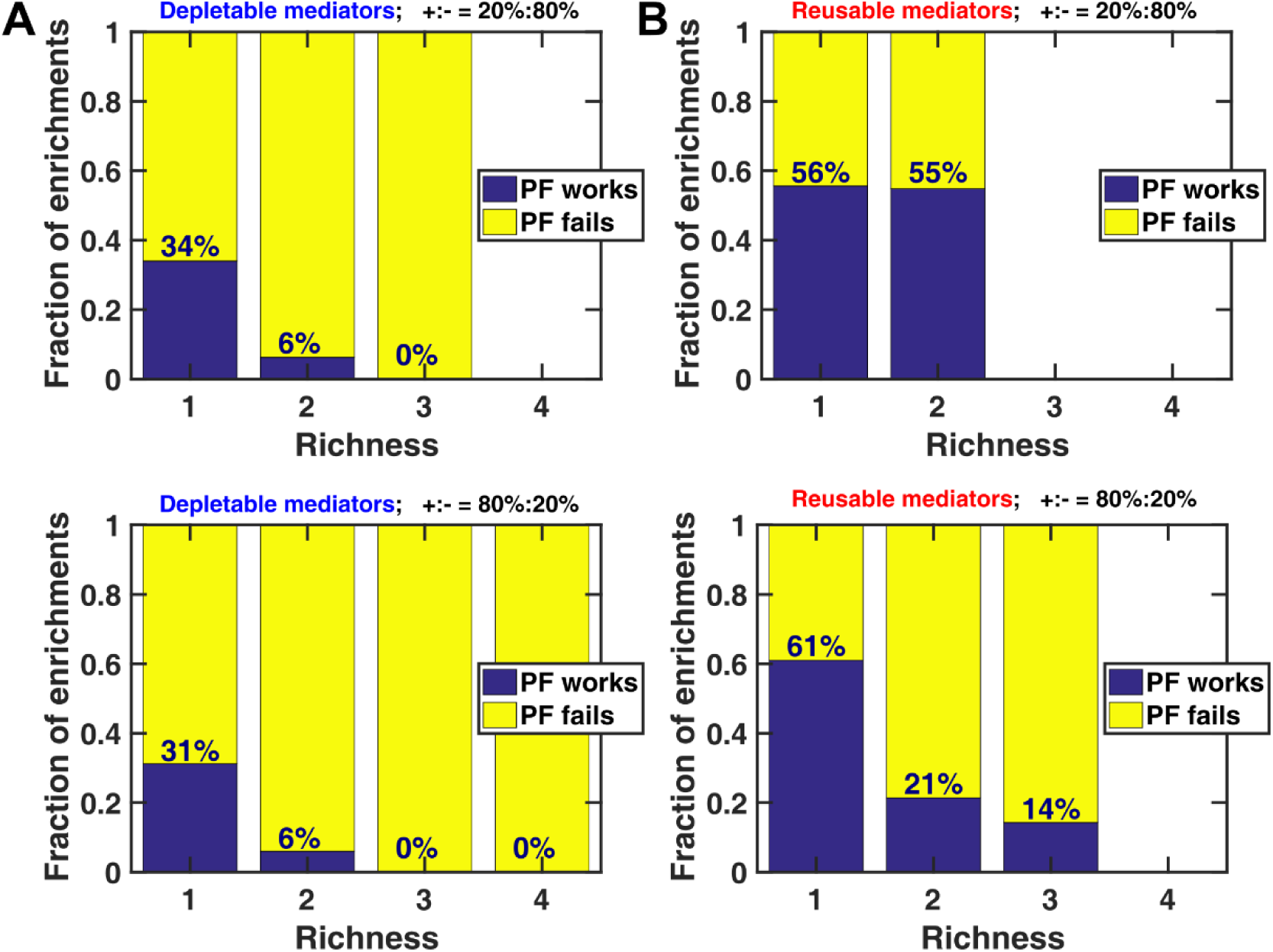
Pairwise model is rarely successful in predicting coexistence of species interacting through chemical mediators. We investigated whether the pairwise model derived for the initial pool of microbes predicts the same species coexist as the reference mechanistic model in more cooperative (+) or competitive (-) environments. Our results suggest that the canonical Lotka-Volterra pairwise model rarely predicts coexisting species in communities with only depletable mediators (A) regardless of the ratio of positive to negative influences (top versus bottom). In communities with only reusable mediators, pairwise models perform better than depletable mediators (i.e. had predictions closer to the reference model), but still fail to reliably predict coexisting species (B). In these simulations, the initial number of species types N_c_=7 and the number of possible mediators N_m_=4. The number of communities analyzed N_s_=1000.

**Fig 4-FS2.**
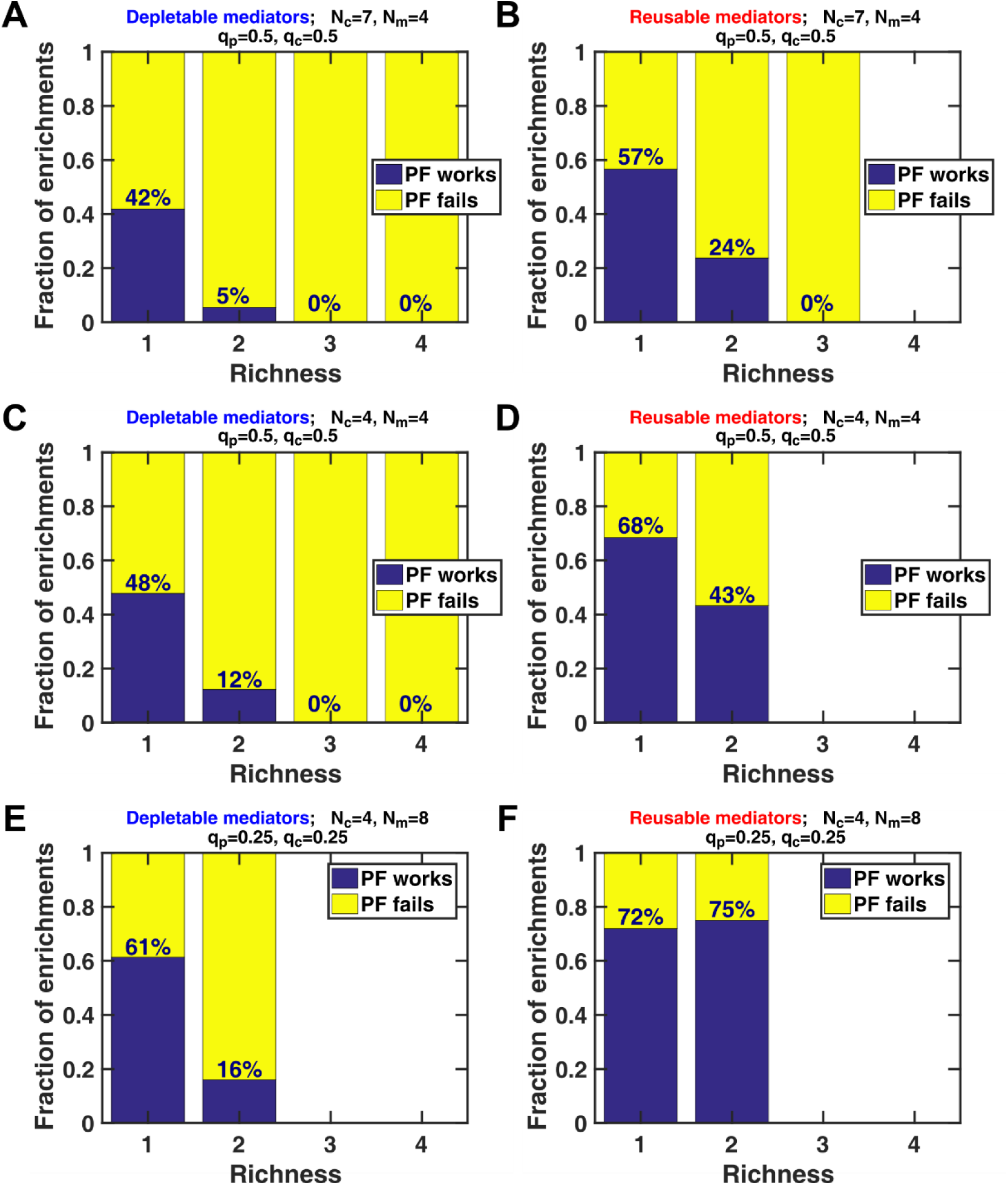
Pairwise model is more appropriate for communities in which interactions are independent. We investigated whether the pairwise model derived for the initial pool of microbes predicts the same species to coexist as the reference mechanistic model when interactions are more independent (i.e. when they take place through independent chemical mediators). In this situation, since mediators are less likely to be shared, the impact of higher-order interactions is reduced. We show three examples: In (A-B), there are seven species in the initial pool, interacting through four mediators, with the chance of the presence of production links (***q_p_***) and the chance of the presence of influence links (***q_c_***) both set at 0.5. With the same parameters as (A-B), in (C-D) we reduced the number of species in the initial pool to four, effectively reducing the chance of overlap in chemical production/removal between interacting species. In (E-F), we increased the number of mediators to eight, but reduced ***q_p_*** and ***q_c_*** to 0.25, maintaining the same chance of interactions as (C-D), but making it more likely for those interactions to take place through independent mediators. Our results suggest that the canonical Lotka-Volterra pairwise model is not suitable for predicting coexisting species in communities with only depletable mediators (A, C, and E). For communities with reusable mediators, however, as interactions become more independent (from B to D, to F), pairwise model becomes more successful in predicting coexistence. The number of communities analyzed in each case, N_s_=1000.

**Fig 4-FS3.**
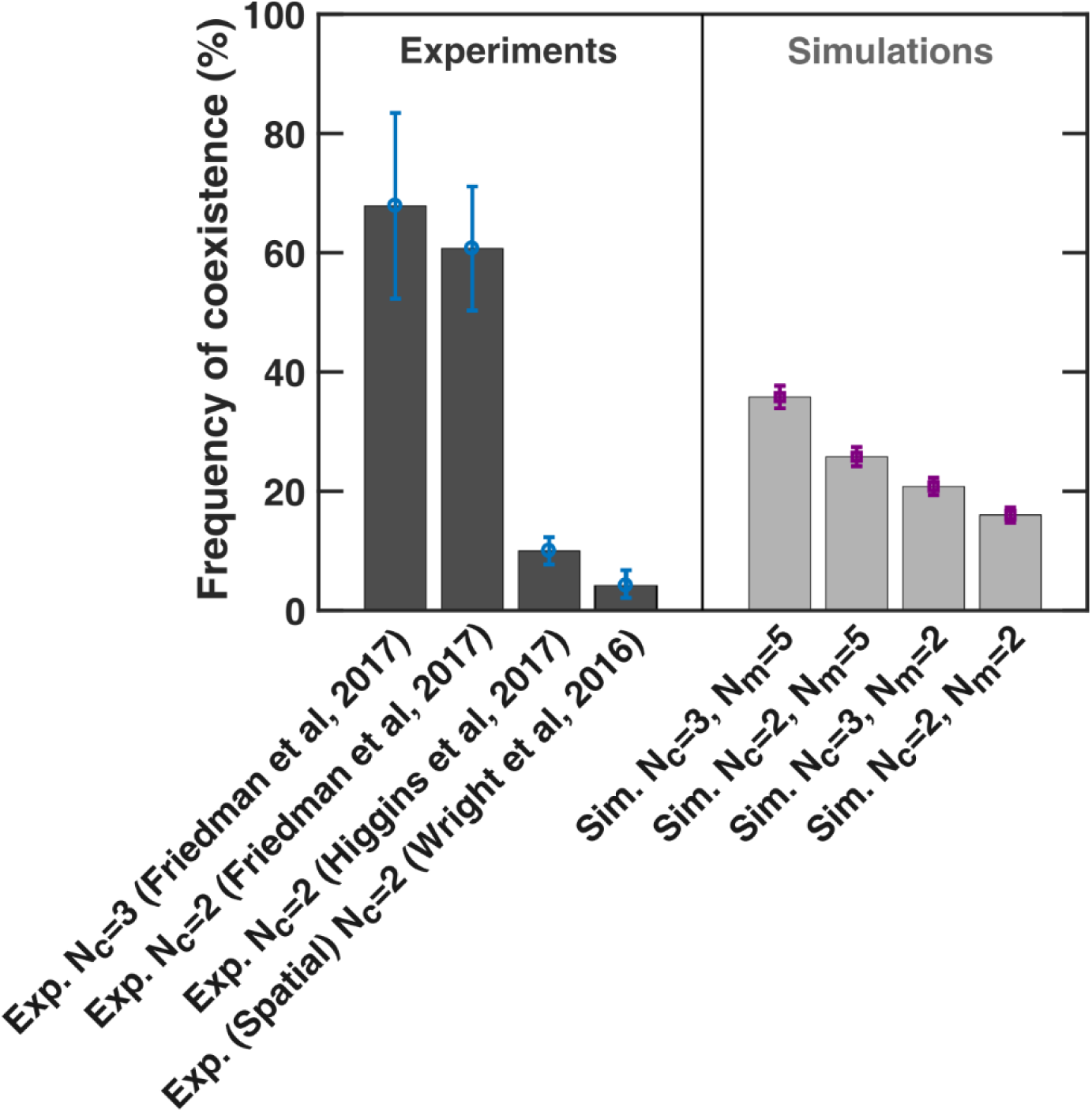
Model predictions cover the range of observed experimental enrichment outcomes. We examined the frequency of achieving coexistence between simulations from our model and reported experimental observations. In simulations, we examined 2- and 3-species initial pool, with 2 or 5 mediators (right, dark gray; simulation parameters are listed in Supplementary Information). Results from available experimental observations come from three reports: Friedman et al investigated coexistence among soil isolates (66); Higgins et al examined a larger set of pairwise interactions among soil isolates in a lab environment (69); and Wright & Vetsigian examined cocultures of pairs of strains from the genus *Streptomyces*, observing mostly invasion and only at a low rate, coexistence (90). Because of the large variability in the experimental data, we speculate that the ecology of strains being examined in each dataset plays a crucial role in their coexistence. We posit that more experimental examples, along with a better mechanistic understanding of interactions in each case is needed to decipher how interactions impact coexistence and if a mediator-explicit model is a suitable representation.

**Fig 5-FS1.**
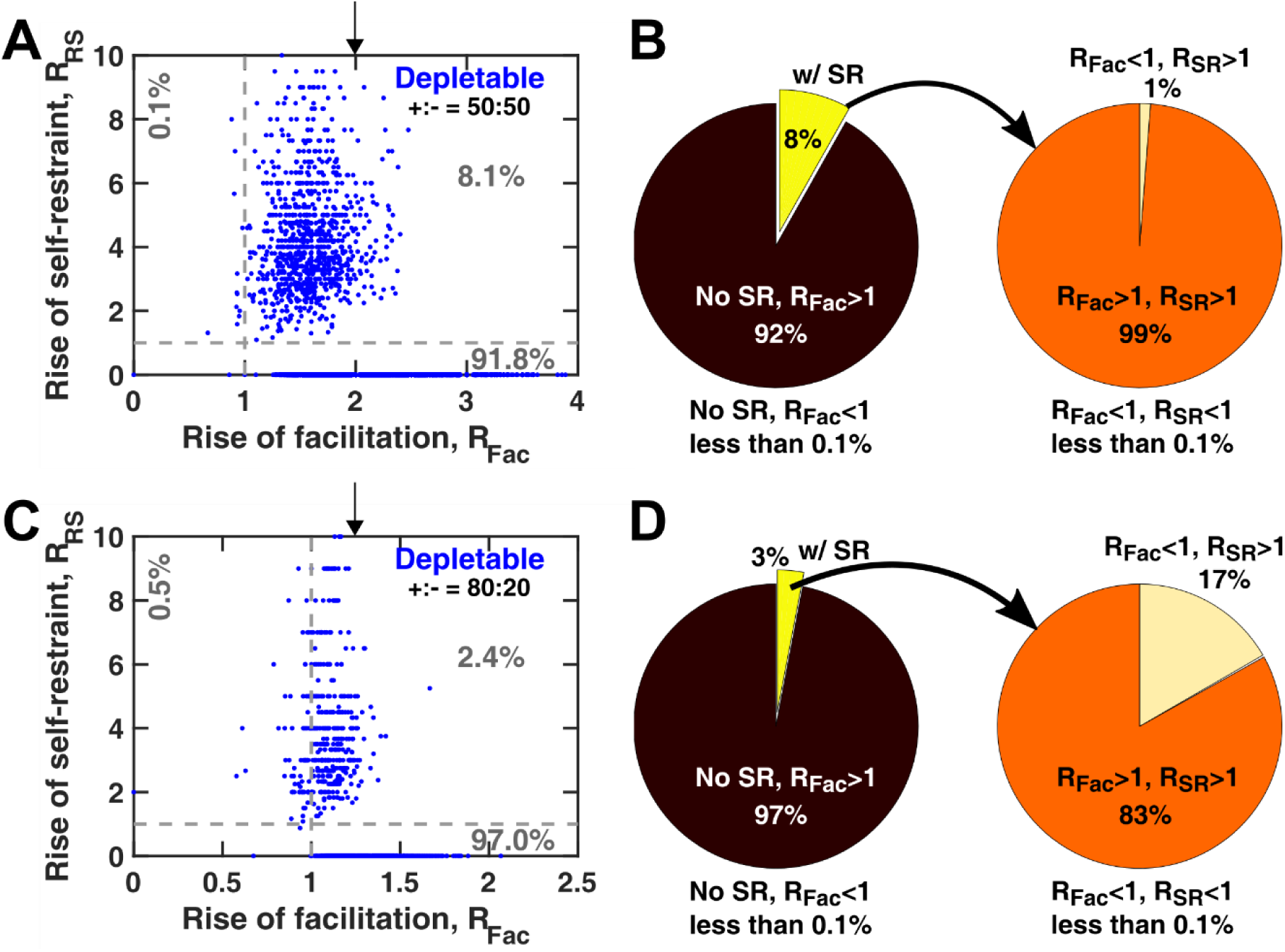
Facilitation and self-restraint are favored during enrichment, regardless of the details of model parameters. Similar to Fig 5, we examined the change in frequencies of facilitation and self-restraint from the initial pool to enriched communities that showed coexistence. Whether the initial pool had equal chance of facilitation versus inhibition (A-B) or was dominated by facilitation (C-D), there was a rise in frequency of facilitation and self-inhibition during enrichment. The arrows in (A) and (C) show the value of rise equal to the inverse of the mean of facilitation frequency in the initial pool. Break-down of different categories in (B) and (D) shows that a rise in facilitation is prevalent, whereas in a small fraction of cases in which facilitation is not dominant a rise in self-inhibition is observed (0.1% in (A), which is 1% of enriched communities that contained self-inhibition (B); 0.5% in (C), which is 17% of enriched communities that contained self-inhibition (D). The number of communities analyzed N_s_=30000. In these simulations, the initial number of species types N_c_=20 and the number of possible mediators N_m_=15. Other simulation parameters are listed in the Supplementary Information (Simulation parameters).

**Fig 6-FS1.**
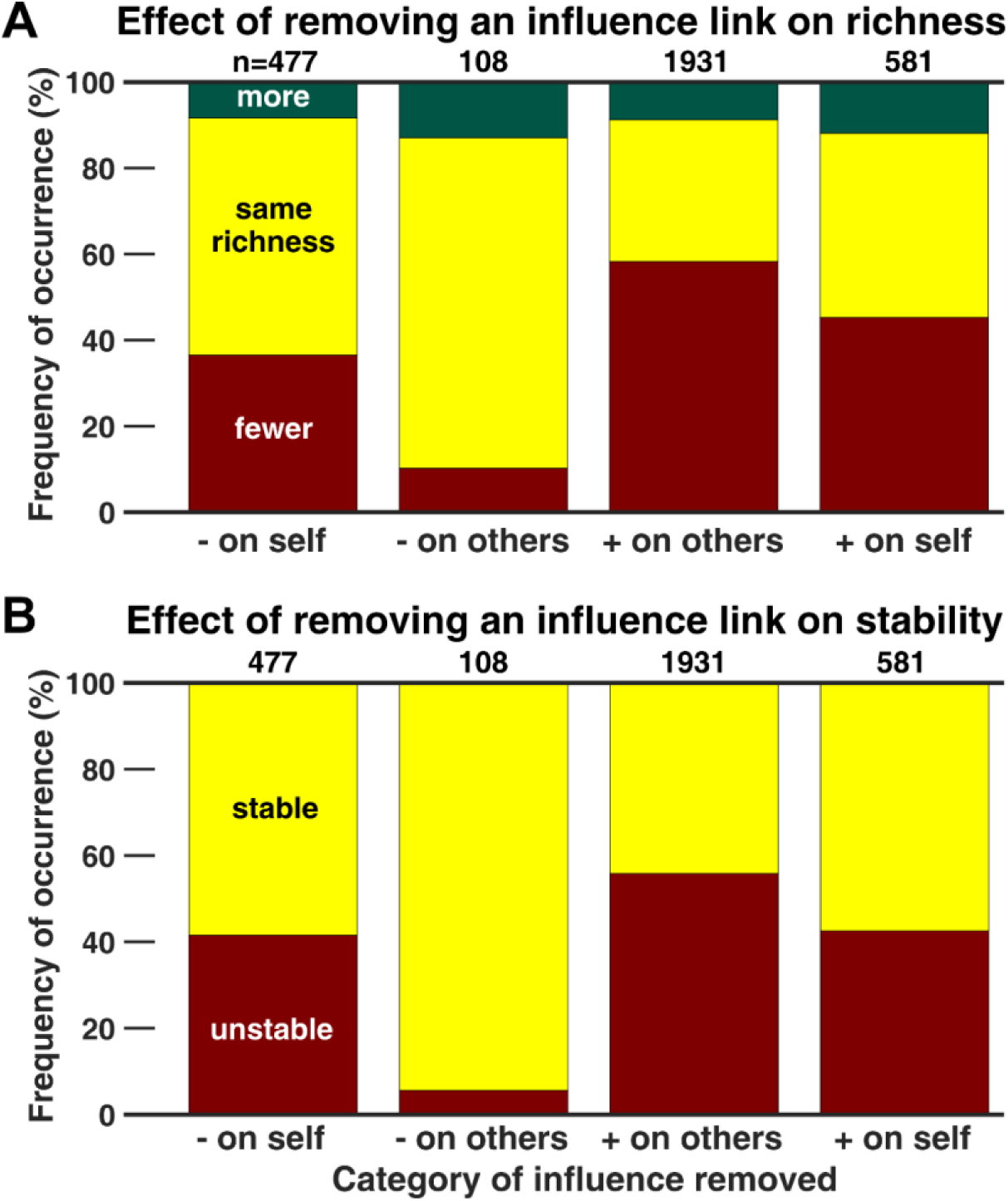
Individual influences can impact coexistence and stability regardless of the interaction makeup in the initial pool. We use knock-out experiments, similar to Fig 6 to assess how removing a link from the interaction network affects coexistence and stability. Here, the initial pool has a binomial network and contains mostly negative influences (+:-= 20%:80%). The conclusions are similar to Fig 6. (A) Removing facilitation links likely disrupts coexistence whereas removing inhibition of others is unlikely to break down coexistence. (B) Removing facilitation likely disrupts the stable community, whereas removal of inhibition of others is unlikely to impact the community.

**Fig 6-FS2.**
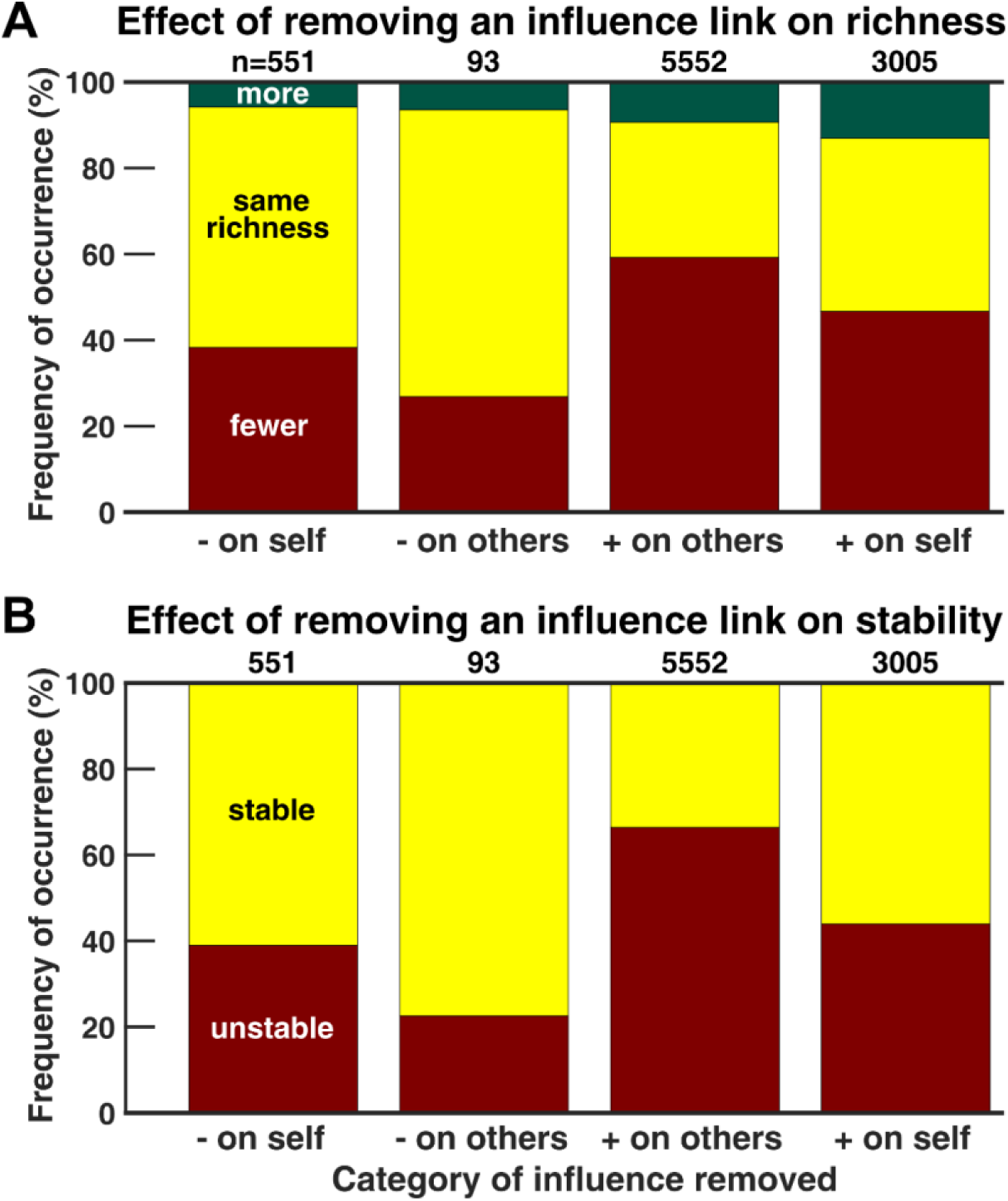
Individual influences can impact coexistence and stability regardless of the interaction network architecture. We use knock-out experiments, similar to Fig 6 to assess how removing a link from the interaction network affects coexistence and stability. Here, the initial pool has a power-law network with average connectivity similar to Fig 6. We largely observe the same conclusions, in which removing facilitation likely disrupts (A) coexistence and (B) stability. Compared to binomial networks in Fig 6, removing inhibition in this case appears more likely to disrupt coexistence and stability, but still not as much as other categories of influences. Similar to Fig 6, the ratio of facilitation to inhibition is chosen to be +:-= 50%:50% in the initial pool for this example.

### Author contribution

Lori Niehaus: ran simulations, analyzed the data, performed the facilitation assessment experiments, and wrote the manuscript
Ian Boland: ran simulations, analyzed the data, and wrote the manuscript
Minghua Liu: developed the code and analyzed the data
Kevin Chen: performed the inhibition assessment experiments
David Fu: performed the inhibition assessment experiments
Catherine Henckel: created the auxotrophic strains and performed the facilitation assessment experiments
Kaitlin Chaung: ran simulations and analyzed the data
Suyen Espinoza: ran simulations and analyzed the data
Samantha Dyckman: created the auxotrophic strains and edited the manuscript
Matthew Crum: ran simulations and analyzed the data
Sandra Dedrick: performed the inhibition assessment experiments and edited the manuscript
Wenying Shou: conceived the idea and edited the manuscript
Babak Momeni: conceived the idea, developed the code, ran simulations, analyzed the data, performed the inhibition assessment experiments, supervised the project, and wrote the manuscript

## Acknowledgements

We would like to thank the Shiaris lab at the University of Massachusetts, Boston for sharing the *E. coli* K12 strain, and Prof. Masaharu Ishii at the University of Tokyo for sharing the *Brevibacillus* strain with us. Research in Momeni Lab is supported by the Start-up fund provided by Boston College, and by an Award for Excellence in Biomedical Research from Smith Family Foundation. BM would like to thank Welkin Johnson and Sarah McMenamin for their support in the process of preparing this manuscript. We are grateful to Jeff Gore for valuable feedback. BM also thanks Boston College for Undergraduate Research Fellowships that supported the participation of undergraduate students in this work. We also thank Boston College’s Research Services for the Linux Cluster that was used for simulations in this manuscript.

